# Highly contiguous genome assemblies of *Photobacterium* strains isolated from fish light organs using nanopore sequencing technology

**DOI:** 10.1101/2022.10.10.511632

**Authors:** AL Gould, JB Henderson

## Abstract

Several species of luminous bacteria in the genus *Photobacterium* are the light organ symbionts of teleost fishes. *Photobacterium leiognathi* and its subspecies, *P. mandapamensis*, in particular, commonly form bioluminescent symbioses with fish hosts in the Leiognathidae and Acropomatidae families as well as with cardinalfish in the genus *Siphamia* (Apogonidae). These two closely related lineages of *Photobacterium* are right at the cutoff average nucleotide identity used to delimit bacterial species (95-96%) and show overlapping ecological niches, including their host fish range. However, there are only a few whole genome assemblies available for these bacterial species, particularly for symbiotic strains isolated from fish light organs, that can be used to explore genome evolution of these two lineages. Here we used Oxford Nanopore Technologies sequencing to produce long reads for assembling highly contiguous genomes of *Photobacterium* strains isolated from fish light organs, including several *P. kishitanii* strains isolated from deep water fishes. We were able to assemble 31 high-quality genomes with near complete BUSCO scores, many at the chromosome-level, and compare their gene contents, including plasmid genes. In doing so, we discovered a new candidate species of *Photobacterium*, Candidatus *Photobacterium acropomis*, which originated from the light organ of the acropomid fish, *Acropoma japonicum*. We also describe a lack of congruency between the presence of the *luxF* gene, which is involved in light production, and the phylogenetic relationships between closely related *P. leiognathi* and *P. mandapamensis* strains. In contrast, there was strong congruency between *luxF* and the host fish family of origin, suggesting this gene might be essential to initiate bioluminescent symbioses with certain hosts, including *Siphamia* and *Acropoma* species. Our study shows the benefit of using long reads in the assembly of bacterial genomes and outlines an assembly pipeline that results in highly contiguous genomes, even from low-coverage ONT reads.

## Introduction

The genus *Photobacterium* belongs to the Vibrionaceae family of bacteria and contains several luminous species that form symbiotic relationships with a range of fish and squid hosts. *Photobacterium leiognathi* and its subspecies, *P. mandapamensis*, both associate with a broad range of teleost fish hosts, including fish in the Leiognathidae and Acropomatidae families as well as cardinalfish in the genus *Siphamia* (Apogonidae) (Kaeding *et al*. 2007). *Photobacterium kishitanii*, on the other hand, is typically found in colder waters and associates with deep-dwelling fish hosts (Ast & Dunlap 2005). Light production in *Photobacterium* is controlled by a contiguous set of genes, termed the *lux-rib* operon. These genes can vary between species and have been used, in combination with certain housekeeping genes, to distinguish between closely related species (e.g. Ast & Dunlap, 2004, 2005, Dunlap & Ast, 2005, Wada *et al*., 2006, Ast *et al*., 2007b, Kaeding *et al*., 2007, Urbanczyk *et al*. 2011a). The *luxF* gene in particular is a key distinguishing feature between *P. leiognathi* and other luminous *Photobacterium* species, including subspecies *P. mandapamensis; luxF* is present in most species but has been secondarily lost in *P. leiognathi* (Ast & Dunlap 2004). In a recent study in which the *lux* operons of *P. mandapamensis* and *P. leiognathi* were cloned into *E. coli* revealed that *luxF* is not required for light production, but cultures harboring the gene emit more light than without *luxF* (Brodl *et al*. 2022). There are two additional genes located upstream of the *lux-rib* operon, *lumP* and *lumQ*, which encode proteins of the lumazine operon (Ast *et al*. 2007) that also vary between *P. leiognathi* and *P. mandapamensis;* like the *luxF* gene, *lumP* is present in *P. mandapamensis* but absent in *P. leiognathi*. Furthermore, two sets of orthologous genes involved secretion have also been shown to be good at discriminating between the two lineages (Urbanczyk *et al*. 2013).

Despite the divergence between *P. leiognathi* and *P. mandapamensis* in certain genes, the two groups remain indistinguishable at the 16S rRNA gene (Ast & Dunlap 2004, Wada *et al*. 2006), and the average nucleotide identity between the two are slightly above the 95% cut-off of the bacterial species definition, indicating the two should be considered the same species (Urbanczyk *et al*. 2013). Furthermore, cardinalfish in the genus *Siphamia* appear to only associate with *P. mandapamensis* (Kaeding *et al*. 2007, Gould *et al*. 2021), indicating there may be important ecological and/or physiological differences between the two groups that are recognizable by *Siphamia* hosts. For example, they differ in their growth and luminescence responses to salinity as well as the color of light produced (Ast and Dunlap 2004). Urbanczyk *et. al*. carried out a whole genome comparison between a single *P. leiognathi* and *P. mandapamensis* strain, and determined that the *P. leiognathi* strain has a larger genome with higher plasticity and a higher rate of foreign gene acquisition compared to the *P. mandapamensis* strain (Urbanczky *et al*. 2013). However, there is currently a limited number of genomes available with which to investigate the breadth of their genomic differences and how these differences may relate to host range and specificity, particularly for the highly specific association between *P. mandapamensis* and *Siphamia* hosts.

There are currently 18 *P. leiognathi* genomes available from NCBI, three of which are additional assemblies of previously assembled genomes. Of the unique strains for which whole genomes are available, eight originated from the light organs of five distinct fish species, only two of which are assembled at the scaffold level; *P. leiognathi* strain lrivu4.1 (GCA_000509205.1) is comprised of 20 scaffolds (Urbanczyk *et al*. 2013) and the *P. mandapamensis* reference strain svers1.1 (GCA_000211495.1) contains 11 scaffolds (Urbanczyk *et al*. 2011b). However, the genome of *P. mandapamensis* isolated from a non-luminous *Loligo* squid from Singapore was recently assembled using Oxford Nanopore Technologies (ONT) sequencing (Soh *et al*. 2018) and is comprised of only three contigs, representing the large and small chromosome present in most vibrio genomes (Okada *et al*. 2005) as well as one plasmid sequence. There are also 24 *P. kishitanii* genomes currently available on NCBI, four of which are scaffolded, and only the reference strain, ANT-2200 (Ali *et al*. 2010), contains fewer than 50 contigs.

In this study we set out to characterize and compare the genome variation in symbiotic *Photobacterium* strains isolated from fish light organs using ONT sequencing. We focused most of our efforts on *P. leiognathi* and *P. mandapamensis* to gain a more complete understanding of the distinction between these two groups and to look for evidence of genomic traits associated with host range. We also include four symbiotic *P. kishitanii* strains isolated from deep sea fishes as a point of comparison to the *P. leiognathi* and *P. mandapamensis* strains and compare them to the currently available *P. kishitanii* genomes. To assemble these genomes from ONT sequences, we performed various assembly pipelines comprised of different combinations of filtering, polishing, and scaffolding steps and present quality assessments of the different approaches implemented. In total we assembled 31 highly contiguous *Photobacterium* genomes, including several that are fully circularized, and present here a more complete picture of the genome biology of symbiotically luminous *Photobacterium* species associated with fish hosts.

## Methods

### Bacterial isolates and DNA extraction

The luminous bacterial strains in this study were initially isolated from the light organs of various fish species listed in Table 1. Several strains were recently isolated from the light organs of *Siphamia tubifer* collected from Verde Island, Philippines, and from Okinawa, Japan (Table 1). Those fish were handled and euthanized using an approved protocol by the Institutional Animal Care and Use Committee at the California Academy of Sciences. The isolates were each grown on LSW-70 (Kaeding *et al*. 2007) agar plates and resuspended in liquid media overnight. Cell pellets were spun down and rinsed with 1x PBS prior to DNA extraction. High molecular weight DNA was then extracted from the fresh cell pellets using a Qiagen MagAttract HWM DNA kit following the manufacturer’s protocol. Following extraction, the DNA was purified with sparQ PureMag Beads (Quantabio) and final DNA concentrations were determined using the Qubit dsDNA HS kit and a Qubit 3.0 fluorimeter (Thermo Fisher).

**Table 1.**
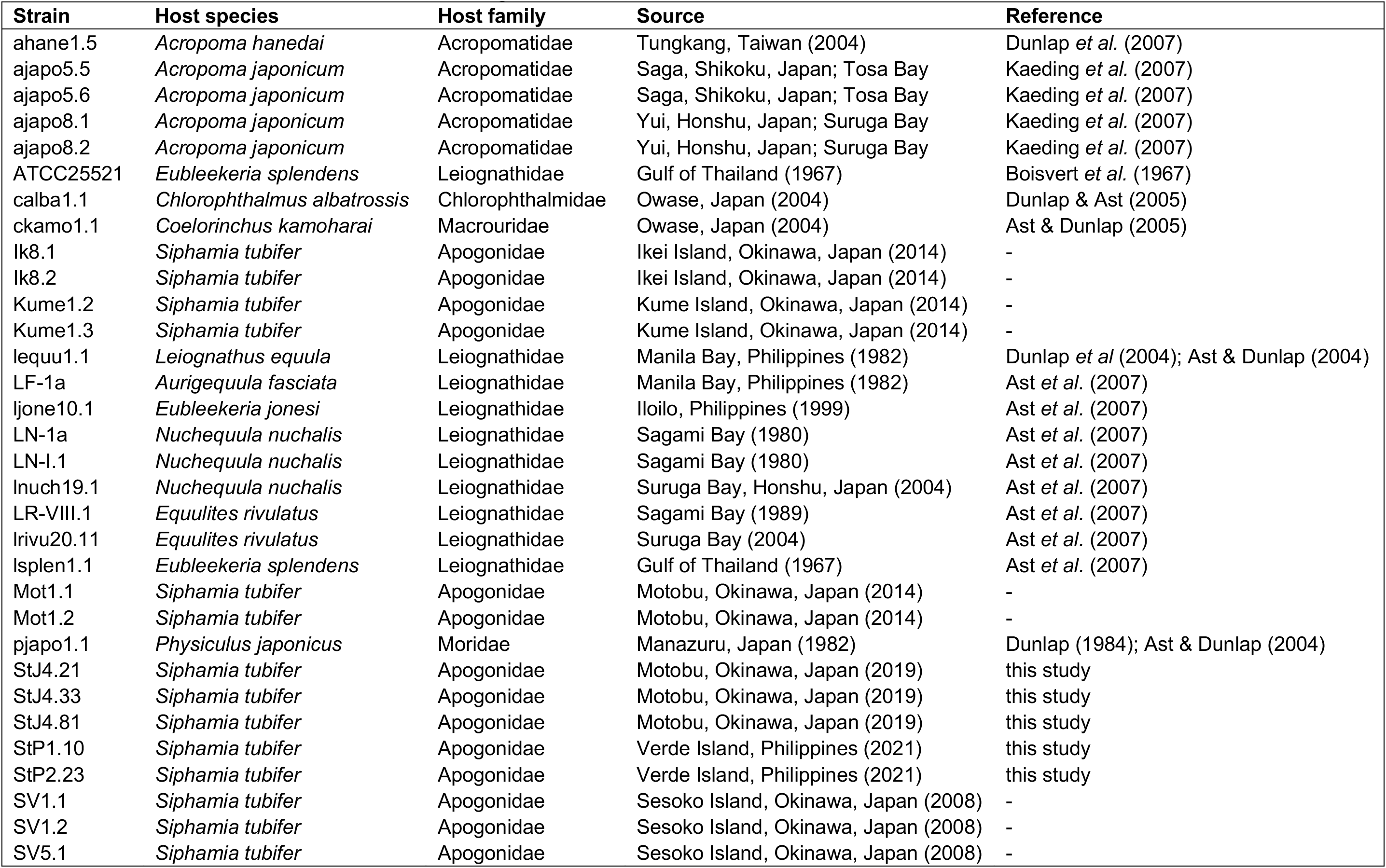
Summary of the 32 *Photobacterium* sp. strains from fish light organs sequenced using Oxford Nanopore Technology. The host species and family from which each strain originated as well as the location and year of collection, when available are listed.

### Library prep and MinION sequencing

DNA concentrations were standardized across samples to an input value of 5.5ng/ul and sequence libraries were prepared with the Rapid (96) Barcoding Kit (Oxford Nanopore Technologies) per the manufacturer’s instructions. The final libraries were pooled and sequenced on a MinION R9.4.1 flow cell. Base-calling for was performed with Guppy v.6.1.7 using the “dna_r9.4.1_450bps_hac” model and a quality score cutoff of eight to retain reads that were used for all subsequent analyses.

### Genome assembly

After base-calling, the sequence reads were additionally filtered with Filtlong (https://github.com/rrwick/Filtlong), removing reads less than 1,000 bp and applying various “keep_percent” settings (80%, 90%, and 95%). Draft genome assemblies were produced from these sets of filtered reads using the Flye assembler (Kolmogorov *et al*. 2019). Circlator (Hunt *et al*. 2015) was then run on the draft assemblies to attempt to circularize any additional contigs, followed by two polishing steps. The first round of polishing was carried out with Medaka (https://github.com/nanoporetech/medaka) followed by Homopolish (Huang *et al*. 2021) with *“Photobacterium”* provided as the input genus. After polishing, additional genome scaffolding was carried out using both RagTag (Algone *et al*. 2021) and Ragout (Kolmogorov *et al*. 2014). The highest quality, circularized and polished draft assemblies produced by Flye were used as references for scaffolding along with the reference strain JS01 described above. For the four *P. kishintanii* strains, the reference genome (ANT-2200, GCA_002631085.1), which has the fewest number of contigs (n=5) of all *P. kishitanii* genomes available fromn NCBI, was used for scaffolding.

### Hybrid assemblies

Two strains, StP2.23 and StJ4.81, also had Illumina short reads (150 bp paired-end reads) available from a recent study (Gould *et al*. in prep) that were used along with the ONT reads as input for Unicycler (Wick *et al*. 2007) to produce hybrid assemblies. The short reads were first quality filtered and trimmed using fastp (Chen *et al*. 2018). The resulting assemblies were also circularized and scaffolded with Circlator (Hunt *et al*. 2015) and RagTag (Algone *et al*. 2021), respectively, and compared with their long read-only assemblies.

### Annotation and genome comparisons

BUSCO (Seppey *et al*. 2019) scores were calculated using the Vibrionales (vibrionales_odb10) set of genes (n=1,445) throughout the assembly pipeline to assess completeness. Similarly, Prokka (Seemann 2014) was implemented to annotate the draft assemblies at each step and to compare gene content and number. QUAST (Gurevich *et al*. 2013) was used to calculate genome statistics at various steps as well.

### Pangenome and phylogenetic analysis

A pangenome analysis of the *P. leiognathi* and *P. mandapamensis* strains was carried out with Roary (Page *et al*. 2015) based on the Prokka annotations of the final assemblies. Additional reference strains available from NCBI (Table 2) were included in the analysis for comparison. The core alignment produced by Roary was then used to construct a maximum likelihood phylogeny in IQ-TREE (Nguyen *et al*. 2015) using the best predicted model (GTR+F+I+G4) and a maximum of 1,000 bootstrap replicates. An additional pangenome analysis was carried out with Roary on all strains, including the four *P. kishitanii* strains, and a phylogeny including these strains was inferred from the core genome alignment with IQ-TREE as previously described. Whole genome comparisons were made between all pairwise combinations of strains using FastANI (Jain *et al*. 2018), and ANIclustermap v1.2.0 (Shimoyama 2022) was implemented to visualize the results.

**Table 2.**
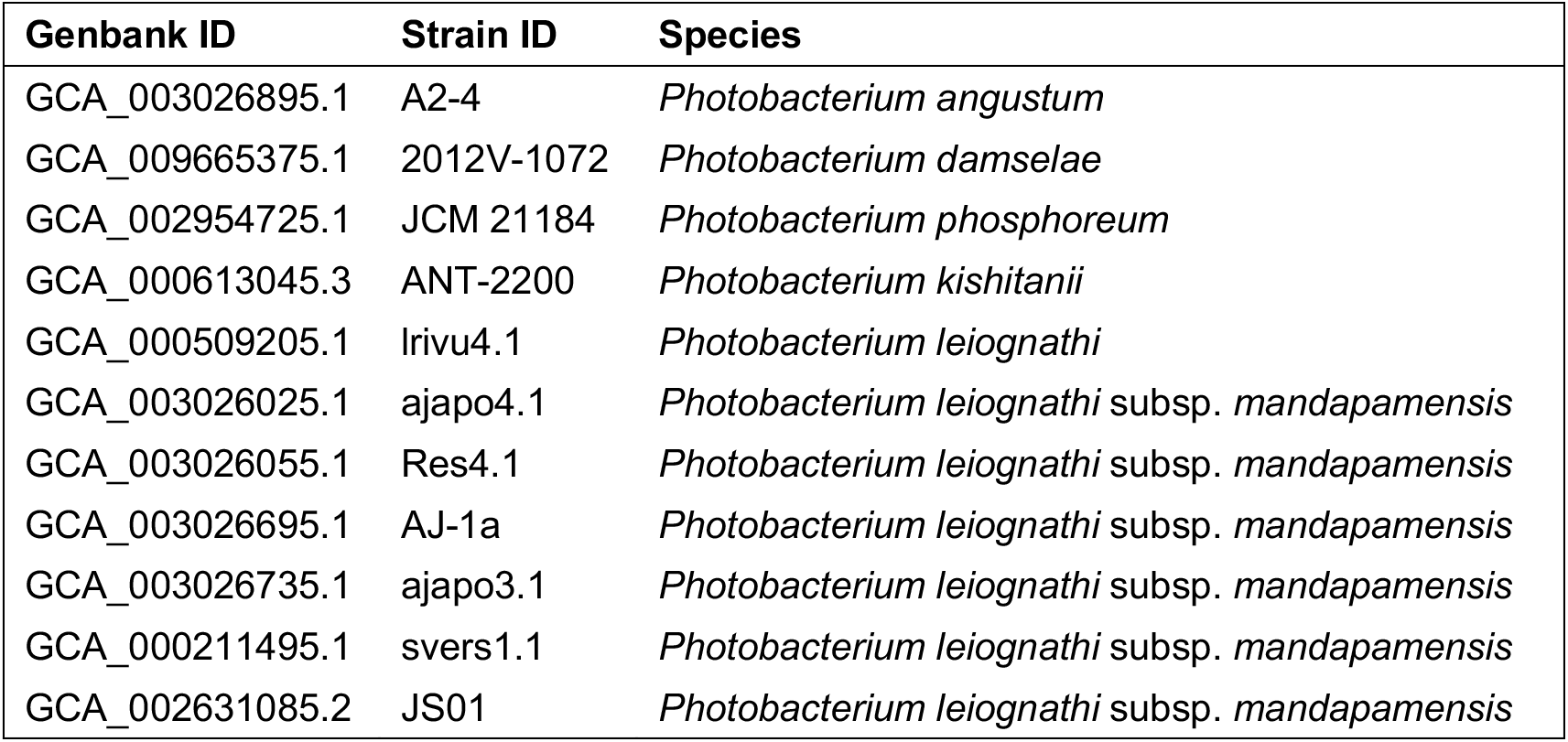
*Photobacterium* genomes available from NCBI included in the study.

### Plasmids

Plasmids from 106 Photobacterium bacteria downloaded from GenBank July 26, 2022 together with the *P. mandapamensis* strain Ikei8.2 contig manually determined to be its plasmid from this study were combined and used to create a blast nucleotide database (makeblastdbs -in plasmids.fasta -parse_seqids -dbtype nucl). blastn was then run over this database with the final assemblies as the query, excluding contigs greater than 50,000 bp prefixing contig names with organism identifiers. Output with qcovus greater than 10% was retained. Records matching the tophit blastn contig of each assembly were collected and manually curated to identify candidate plasmid sequences. Prokka (Seemann 2014) was the implemented on these sequences to obtain the plasmid gene content for each strain.

## Results

### MinION Sequencing Data

The ONT MinION sequencing run generated 5.29M fast5 reads with an N50 of 9.1Kb. After demultiplexing and base calling with Guppy, a total of 1.76M reads (4.82 Gbp) were obtained, 145,744 of which were unclassified (no barcode could be assigned). The number of reads assigned to each sample ranged from 7,816 to 122,045, and the minimum and maximum sequencing depths were 4× and 130×, respectively.

### Draft assembly statistics

After the initial Flye assembly on the filtered long reads (90 “keep percent”), one genome, ajapo5.5, was assembled as two complete circular chromosomes, one large (>3,100,000 bp) and one small (>1,400,00 bp) with one additional circular plasmid (~16,000 bp) (Fig. 1). Nine additional genomes assemblies contained at least one circular chromosome (Fig. 1). After running circlator on the Flye assemblies, two additional chromosomes were circularized from two different assemblies. The total number of contigs also decreased for nearly all of the assemblies (Fig. 2). In contrast, the polishing steps had no effect on the number of contigs, but did increase the BUSCO completeness scores, in some cases by a large percentage. Running Homopolish after the initial polishing with medaka especially improved BUSCO scores across all strains, and in some cases, there was a greater than 20% increase in completeness (Fig. 3). Scaffolding, on the other hand, had little to no effect on the BUSCO scores, but did decrease the number of contigs even further nearly all assemblies; 16 of the 32 strains ended up with draft genomes that were comprised of only 2 or 3 contigs (Table S1). The most notable scaffolding improvement was observed for strain StJ4.81, which went from 50 to 3 contigs after scaffolding the polished assemblies. Scaffolding also increased the number of coding sequences (CDS) detected for most strains (Table S1). In the case of StJ4.81, the total number of CDS increased from 4,346 to 4,361, while the number of rRNAs and tRNAs remained the same, 63 and 208, respectively. One strain, StJ4.33, had low average coverage (3.4x) and the BUSCO completeness score only reached 2.8%. Thus, it was removed from further analysis. Of the remaining 31 strains, 27 had BUSCO completeness scores of 95% or greater, 15 of which were 99% complete (Fig. 3).

**Figure 1.**
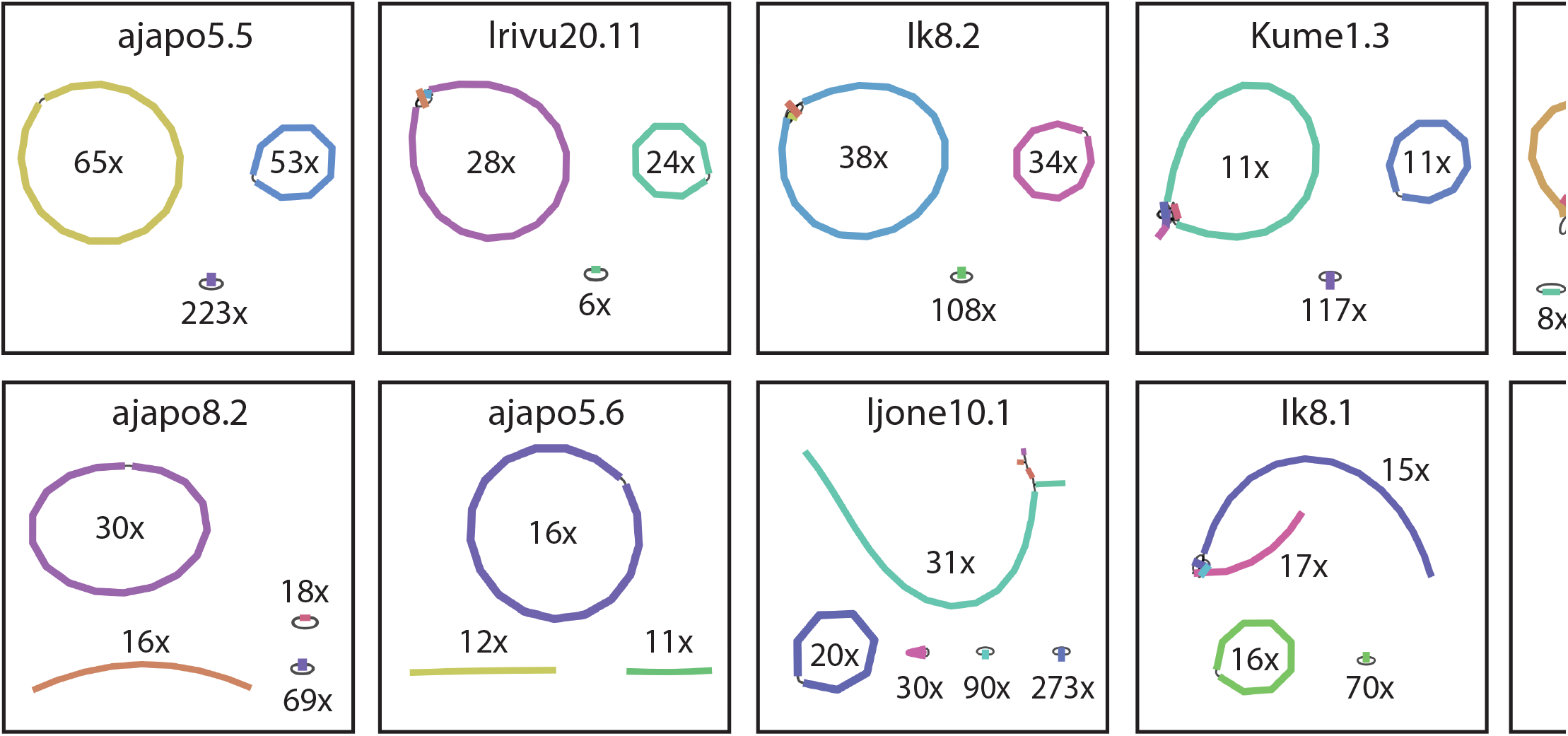
Assembly graphs for the ten *Photobacterium* strains with at least one fully circularized chromosome from the initial Flye assembly. Coverage depth of each contig is also indicated. Plots were made with the program Bandage.

**Figure 2.**
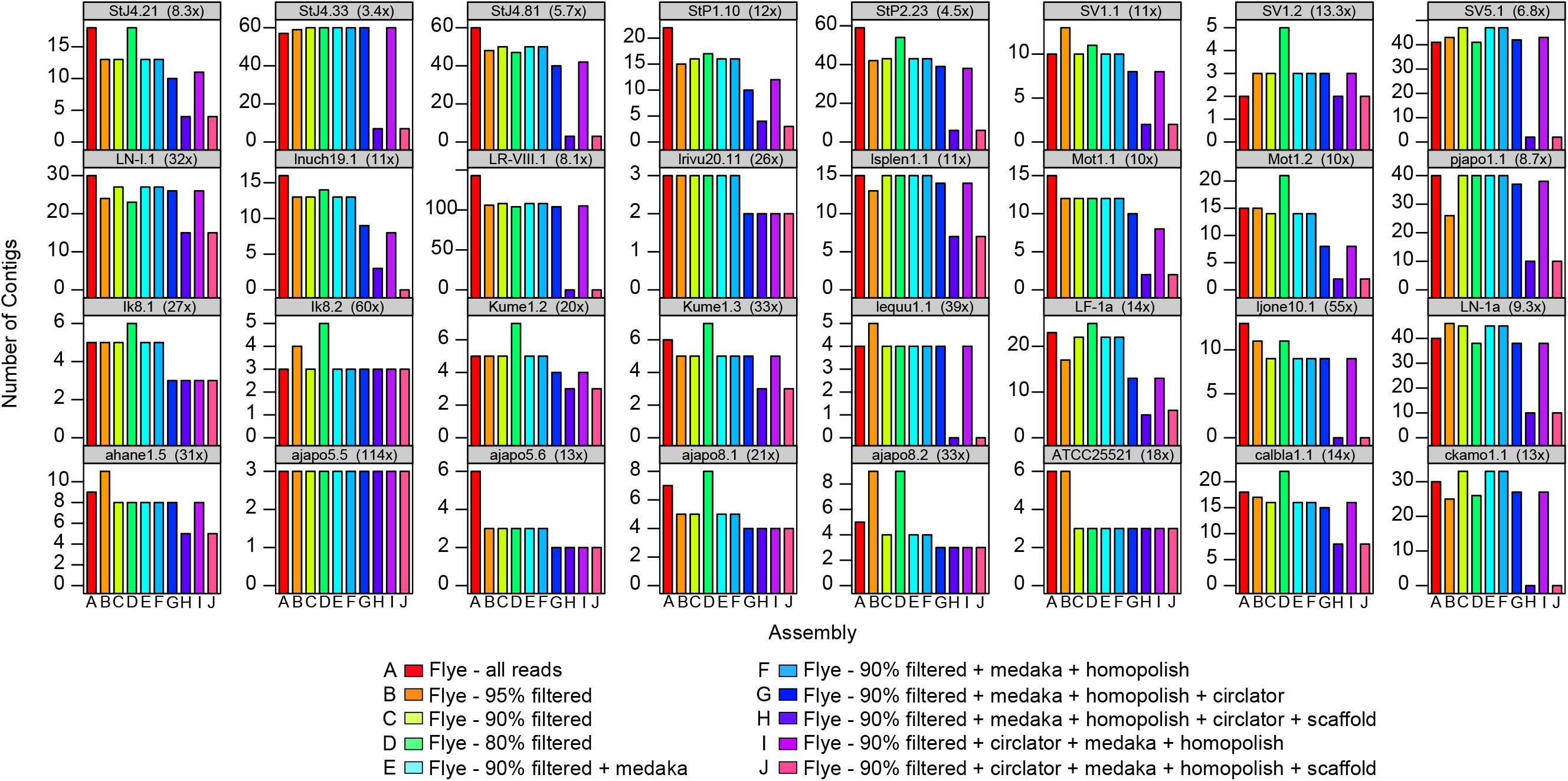
Number of contigs for each draft assembly of the 32 strains of *Photobacterium* using only ONT reads. The strain names and their average coverage depth are indicated in the gray bar above each plot. The different assembly approaches are indicated in the legend and colored accordingly.

**Figure 3.**
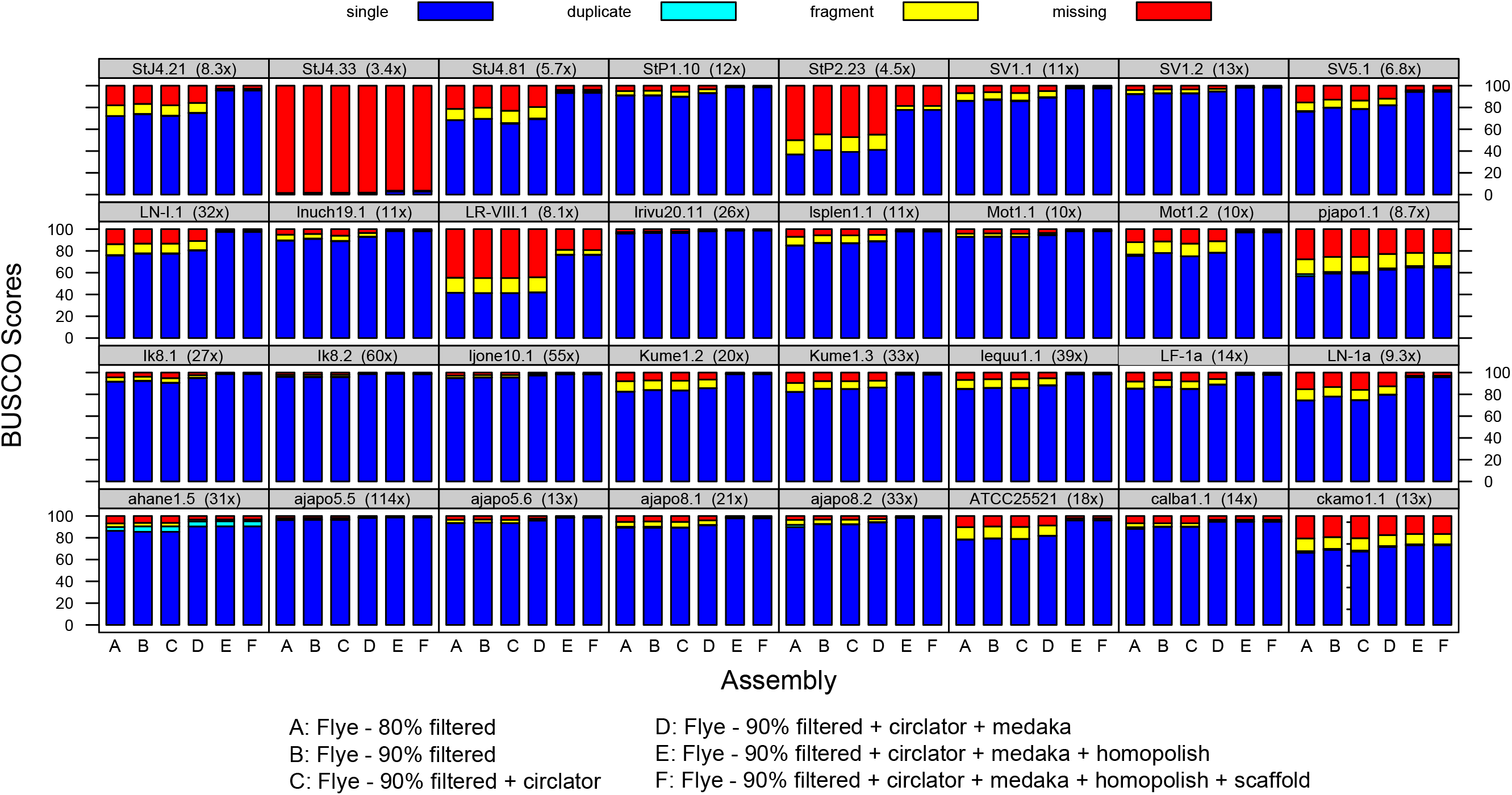
BUSCO scores for the draft assemblies of the 32 strains of *Photobacterium* using only ONT reads. The strain names and their average coverage depth are indicated in the gray bar above each plot. The different assembly approaches are indicated in the legend and the bar colors represent the different BUSCO gene categories: single copy (blue), duplicate (cyan), fragmented (yellow), and missing (red).

For the final assemblies, the average genome size was 4,944,424 bp across all 32 strains, ranging from 4,521,083 to 5,791,416 bp, including four *P. kishitanii* strains. The *Photobacterium leiognathi* and *P. mandapamensis* strains averaged 4,920,253 total bp and had an average of 4,393 CDS, 49 rRNAs, and 193 tRNAs. The *P. leiognathi* strains were approximately 6% larger than the *P. mandapamensis* genomes, whereas the four *P. kishitanii* strains were even larger, averaging 5,085,414 total bp with an average of 5,576 CDSs, 33 rRNAs, and 196 tRNAs (Table 3). For the *P. leiognathi* and *P. mandapamensis* strains with fully circularized chromosomes, the larger chromosome averaged 3,207,570 bp and the smaller one averaged 1,511,137 bp. The assembly for strain Ik8.2, which was isolated from a *Siphamia tubifer* light organ from Okinawa, Japan in 2014, consisted of three circular contigs, representing both chromosomes and a plasmid. There were 2,741 CDSs, 181 tRNAs, and 56 rRNAs on the large chromosome and 1,320 CDSs, 27 tRNAs, and no rRNAs on the small chromosome. Similarly, strain ajapo5.5, which had the highest depth of coverage of all strains, contained 2,696 CDSs, 181 tRNAs, and 59 rRNAs on the large chromosome and 1,286 CDSs, 27 tRNAs, and no rRNAs on the small chromosome. A BLAST comparison of *P. mandapamensis* strain Ik8.2 to several other strains revealed multiple unique gene regions that were only found in the Ik8.2 genome (Fig.4), but most of the genes in these regions were of unknown function. A comparison of the genome assembly of strain Ik8.2 and the reference *P. mandapamensis* strain svers1.1, indicates a high degree of genome synteny and exemplifies the ability of this highly contiguous assembly to be used to scaffold previous assemblies available from NCBI (Fig. 4).

**Figure 4.**
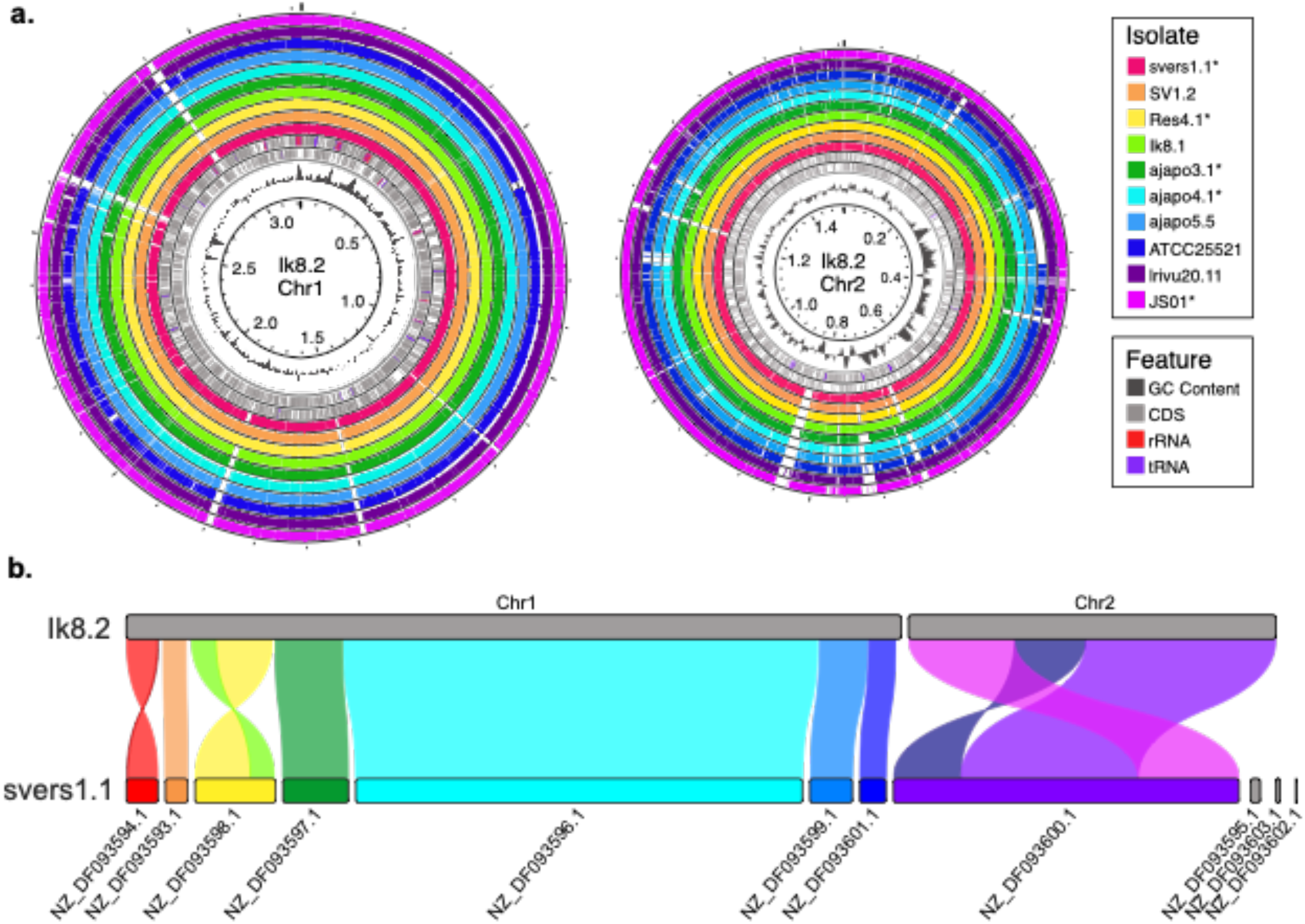
Synteny of the genome assembly of isolate Ik8.2 to that of other *P. leiognathi (mandapamensis*) genomes. a) blastn alignments of the isolates listed to both chromosomes of Ik8.2. Assemblies from NCBI are indicated with an *. Genomic features including coding sequences (CDS), rRNAs, tRNAs, and GC content are shown on the inner rings of each chromosome. b) Syntenic blocks between the *P. leiognathi* subsp. *mandapamensis* reference strain svers1.1 and the chromosome-level assembly of Ik8.2.

**Table 3.** Statistics for the final genome assemblies as determined by QUAST. Reference strains indicated in bold with an * are also included for comparison. Listed are the number of total contigs, the number of contigs greater than 1,000, 10,000, and 50,000 bp, the largest contig in bp, the total number of bp, %GC content, the N50 and L50 values, and the number of N’s per 100 Kbp. The first five shaded entries are *Photobacterium kishitanii* strains whereas all others are *P. leiognathi* and *P. mandapamensis* strains.

| Strain | Total bp | Contigs | Largest contig | N50 | L50 | GC(%) | Ns | CDS | rRNA | tRNA |
| --- | --- | --- | --- | --- | --- | --- | --- | --- | --- | --- |
| ahane1.5 | 5,100,844 | 5 | 3,353,657 | 3,353,657 | 1 | 39.02 | 126 | 4,680 | 53 | 207 |
| calba1.1 | 5,200,973 | 7 | 3,263,220 | 3,263,220 | 1 | 38.96 | 792 | 4,892 | 46 | 206 |
| ckamo1.1 | 5,039,339 | 4 | 3,317,370 | 3,317,370 | 1 | 38.96 | 2,489 | 6,136 | 8 | 173 |
| pjapo1.1 | 5,000,498 | 4 | 3,339,696 | 3,339,696 | 1 | 39.36 | 12,224 | 6,596 | 25 | 196 |
| **pjapo1.1\*** | 4,695,065 | 117 | 925,439 | 174,214 | 6 | 39.10 | 0 | 4,217 | 6 | 84 |
| **AJ-1a\*** | 4,711,244 | 65 | 468,721 | 238,774 | 7 | 41.14 | 0 | 4,156 | 13 | 131 |
| **ajapo3.1\*** | 4,794,394 | 51 | 480,625 | 245,626 | 8 | 40.98 | 0 | 4,214 | 12 | 137 |
| **ajapo4.1\*** | 4,576,643 | 52 | 696,744 | 262,428 | 5 | 41.2 | 0 | 3,991 | 11 | 124 |
| ajapo5.5 | 4,690,822 | 3 | 3,177,212 | 3,177,212 | 1 | 41.37 | 0 | 4,004 | 59 | 307 |
| ajapo5.6 | 4,729,792 | 2 | 3,174,233 | 3,174,233 | 1 | 41.27 | 0 | 4,052 | 62 | 213 |
| ajapo8.1 | 4,878,649 | 4 | 3,245,602 | 3,245,602 | 1 | 41.12 | 0 | 4,255 | 44 | 192 |
| ajapo8.2 | 4,886,267 | 3 | 3,261,968 | 3,261,968 | 1 | 41.15 | 0 | 4,239 | 53 | 194 |
| ATCC25521 | 4,750,881 | 3 | 3,269,131 | 3,269,131 | 1 | 41.02 | 0 | 4,255 | 41 | 194 |
| lk8.1 | 4,765,281 | 3 | 3,231,342 | 3,231,342 | 1 | 41.15 | 0 | 4,130 | 47 | 197 |
| lk8.2 | 4,783,140 | 3 | 3,218,875 | 3,218,875 | 1 | 41.17 | 0 | 4,104 | 56 | 207 |
| **JS01\*** | 4,874,529 | 3 | 3,251,164 | 3,251,164 | 1 | 41.20 | 0 | 4,288 | 57 | 205 |
| Kume1.2 | 4,835,311 | 3 | 3,186,767 | 3,186,767 | 1 | 41.25 | 1,634 | 4,156 | 49 | 197 |
| Kume1.3 | 4,818,955 | 3 | 3,185,044 | 3,185,044 | 1 | 41.23 | 1,911 | 4,137 | 43 | 194 |
| lequu1.1 | 4,825,624 | 4 | 3,204,041 | 3,204,041 | 1 | 41.19 | 0 | 4,192 | 62 | 209 |
| LF-1a | 5,104,659 | 5 | 3,267,821 | 3,267,821 | 1 | 40.96 | 1,222 | 4,607 | 39 | 192 |
| ljone10.1 | 5,276,714 | 8 | 3,313,411 | 3,313,411 | 1 | 41.34 | 1 | 4,535 | 62 | 207 |
| LN-1a | 5,174,290 | 10 | 3,359,026 | 3,359,026 | 1 | 41.21 | 54 | 4,760 | 44 | 195 |
| LN-I.1 | 5,490,629 | 15 | 3,405,483 | 3,405,483 | 1 | 41.53 | 20 | 5,017 | 62 | 200 |
| Inuch19.1 | 5,237,683 | 5 | 3,439,577 | 3,439,577 | 1 | 41.18 | 162 | 4,618 | 57 | 197 |
| LR-VIII.1 | 5,791,416 | 21 | 3,699,906 | 3,699,906 | 1 | 41.28 | 22,860 | 5,602 | 32 | 147 |
| Irivu20.11 | 4,738,701 | 2 | 3,228,638 | 3,228,638 | 1 | 41.24 | 0 | 4,069 | 58 | 209 |
| **Irivu4.1\*** | 5,268,214 | 20 | 1,730,671 | 979,827 | 2 | 40.98 | 6,943 | 4,332 | 3 | 72 |
| Isplen1.1 | 5,292,468 | 7 | 3,284,001 | 3,284,001 | 1 | 41.01 | 813 | 4,768 | 37 | 174 |
| Mot1.1 | 4,676,757 | 2 | 3,973,505 | 3,973,505 | 1 | 41.25 | 13 | 4,069 | 62 | 208 |
| Mot1.2 | 4,947,312 | 2 | 3,234,211 | 3,234,211 | 1 | 41.2 | 2,558 | 4,336 | 62 | 206 |
| **Res4.1\*** | 4,730,847 | 65 | 766,940 | 129,576 | 9 | 40.99 | 0 | 4,098 | 13 | 152 |
| StJ4.21 | 4,877,430 | 4 | 3,228,436 | 3,228,436 | 1 | 41.23 | 623 | 4,361 | 63 | 208 |
| StJ4.81 | 4,713,802 | 3 | 3,157,280 | 3,157,280 | 1 | 41.22 | 6 | 4,047 | 45 | 195 |
| StP1.10 | 4,781,050 | 3 | 3,175,608 | 3,175,608 | 1 | 41.15 | 19 | 4,194 | 47 | 194 |
| StP2.23 | 4,685,648 | 3 | 3,130,376 | 3,130,376 | 1 | 41.17 | 11 | 4,043 | 33 | 182 |
| SV1.1 | 4,749,844 | 2 | 3,174,417 | 3,174,417 | 1 | 41.13 | 1,321 | 4,126 | 38 | 190 |
| SV1.2 | 4,718,353 | 2 | 3,184,478 | 3,184,478 | 1 | 41.17 | 0 | 4,083 | 51 | 200 |
| SV5.1 | 4,521,083 | 2 | 3,089,627 | 3,089,627 | 1 | 41.27 | 91 | 4,122 | 35 | 193 |
| **svers1.1\*** | 4,598,918 | 11 | 1,910,320 | 1,477,894 | 2 | 41.06 | 742 | 4,031 | 6 | 75 |

### Hybrid assemblies

The use of short reads improved the assembly for the two strains for which short reads were available, StP2.23 and StJ4.81. After trimming, there were 3,385,214 and 5,808,314 paired end reads for StP2.23 and StJ4.81, respectively that were used for the initial assembly step in Unicycler (Wick *et al*. 2007). With respect to BUSCO scores, the hybrid assemblies were more complete than the long read-only assemblies. For strain StP2.23, the BUSCO completeness score went from 77.8% to 99.1% for the Flye and Unicycler assemblies, respectively, and for StJ4.81, it improved from 94.5% to 99.1% (Table 4). Running both Circlator (Hunt *et al*. 2015) and RagTag (Algone *et al*. 2021) on the assemblies reduced the number of contigs but had slightly negative effects on the BUSCO scores. The hybrid assembly for StP2.23 went from 30 contigs down to 2 scaffolds, but the BUSCO completeness score decreased to 96.3%. Similarly, the hybrid assembly for strain StJ4.81 went from 15 to 2 contigs after both circularizing and scaffolding, but BUSCO completeness dropped to 98.6% (Table 4). However, removing sequences less than 1,000 bp from the scaffolded (non-circularized) hybrid assemblies resulted in only three contigs for both strains and 99.1% BUSCO completeness scores, and were used in the remaining analyses for strains StP2.23 and StJ4.81.

**Table 4.** Comparison of the long read-only and hybrid draft assemblies for the two *P. mandapamensis* strains with short reads available. Listed are the number of contigs (Ctgs), the total number of bp in the assembly, the largest contig size (bp), the GC content, N50 and L50 values, the average number of Ns/100 kbp, the number of coding sequences (CDS), the number of rRNAs and tRNAs, the number of repeat regions, and the BUSCO scores shown as percentages: complete [single, duplicate], fragmented, and missing.

| Strain | Assembly | Ctgs | Total bp | GC% | N50 | L50 | Ns | CDS | rRNA | tRNA | Repeat | Complete [S, D] | Fragment | Missing |
| --- | --- | --- | --- | --- | --- | --- | --- | --- | --- | --- | --- | --- | --- | --- |
| StP2.23 | flye+circ+polish | 38 | 4228014 | 41.09 | 209731 | 7 | 0 | 4211 | 20 | 144 | 0 | 77.8 [77.7, 0.1] | 3.60 | 18.60 |
|  | flye+circ+polish+scaffold | 4 | 4888897 | 41.09 | 3236904 | 1 | 14038 | 4525 | 20 | 144 | 0 | 77.8 [77.7, 0.1] | 3.60 | 18.60 |
|  | unicycler | 30 | 4686987 | 41.18 | 2741469 | 1 | 0 | 4044 | 33 | 188 | 2 | 99.1 [98.8, 0.3] | 0.10 | 0.80 |
|  | unicycler+circ | 4 | 4688278 | 41.18 | 3000960 | 1 | 0 | 4112 | 35 | 182 | 2 | 96.0 [95.7, 0.3] | 1.20 | 2.80 |
|  | unicycler+circ+scaffold | 2 | 4688478 | 41.18 | 3134673 | 1 | 4.27 | 4112 | 35 | 182 | 2 | 96.0 [95.7, 0.3] | 1.20 | 2.80 |
|  | unicycler+scaffold | 25 | 4691567 | 41.18 | 3130376 | 1 | 10.67 | 4044 | 33 | 188 | 2 | 98.5 [98.0, 0.5] | 0.20 | 1.30 |
|  | **unicycler+scaffold+circ** | **2** | **4689350** | **41.18** | **3135545** | **1** | **4.26** | **4095** | **35** | **181** | **2** | **96.3 [96.0, 0.3]** | **1.00** | **2.70** |
| StJ4.81 | flye+circ+polish | 42 | 4610923 | 41.33 | 215021 | 6 | 0 | 4280 | 46 | 187 | 0 | 94.5 [93.3, 1.2] | 1.60 | 3.90 |
|  | flye+circ+polish+scaffold | 3 | 4614823 | 41.33 | 3008273 | 1 | 84.51 | 4279 | 46 | 187 | 0 | 94.6 [93.4, 1.2] | 1.60 | 3.80 |
|  | unicycler | 15 | 4716676 | 41.23 | 1555055 | 2 | 0 | 4047 | 45 | 196 | 2 | 99.1 [98.8, 0.3] | 0.10 | 0.80 |
|  | unicycler+circ | 2 | 4716071 | 41.23 | 3161731 | 1 | 0 | 4160 | 47 | 198 | 2 | 95.9 [95.6, 0.3] | 1.00 | 3.10 |
|  | unicycler+circ+scaffold | 2 | 4716071 | 41.23 | 3161731 | 1 | 0 | 4161 | 47 | 198 | 2 | 95.9 [95.6, 0.3] | 1.00 | 3.10 |
|  | unicycler+scaffold | 12 | 4716976 | 41.23 | 3157280 | 1 | 6.36 | 4047 | 45 | 196 | 2 | 99.1 [98.8, 0.3] | 0.10 | 0.80 |
|  | **unicycler+scaffold+circ** | **2** | **4711728** | **41.22** | **3157280** | **1** | **6.37** | **4101** | **44** | **195** | **2** | **98.6 [98.3, 0.3]** | **0.10** | **1.30** |

### Pangenome Analysis

A pangenome analysis revealed a total of 18,142 genes across all *P. leiognathi* and *P. mandapamensis* strains examined in this study. Of these, 2,017 genes are ‘core’ genes shared across at least 95% of the strains and 2,884 are ‘shell’ genes shared across 15-95% of strains. The majority of the genes detected (73%, n=13,241), however, are ‘cloud’ genes present in fewer than 15% of the total strains examined (Fig. 5). A separate analysis of only the *P. leiognathi* strains in this study (n=4) revealed a pangenome of 3,237 genes, whereas the pangenome of the *P. mandapamensis* strains (n=27) contains 2,618 genes. Two strains, ajapo5.5 and the reference strain ajapo4.1 (GCA_003026025.1), both of which originated from the same host species, are divergent from the remaining *P. leiognathi* and *P. mandapamensis* strains and share 765 genes that are not present in any of the other strains examined (Fig. 5). Of these genes, 517 are of unknown function, but of the remaining 248 genes with assigned function, several related to macrolide antibiotics and drug resistance. Additionally, the complete operon for urease production (*ureABCDEFG*) was present in both strains (Table S2).

**Figure 5.**
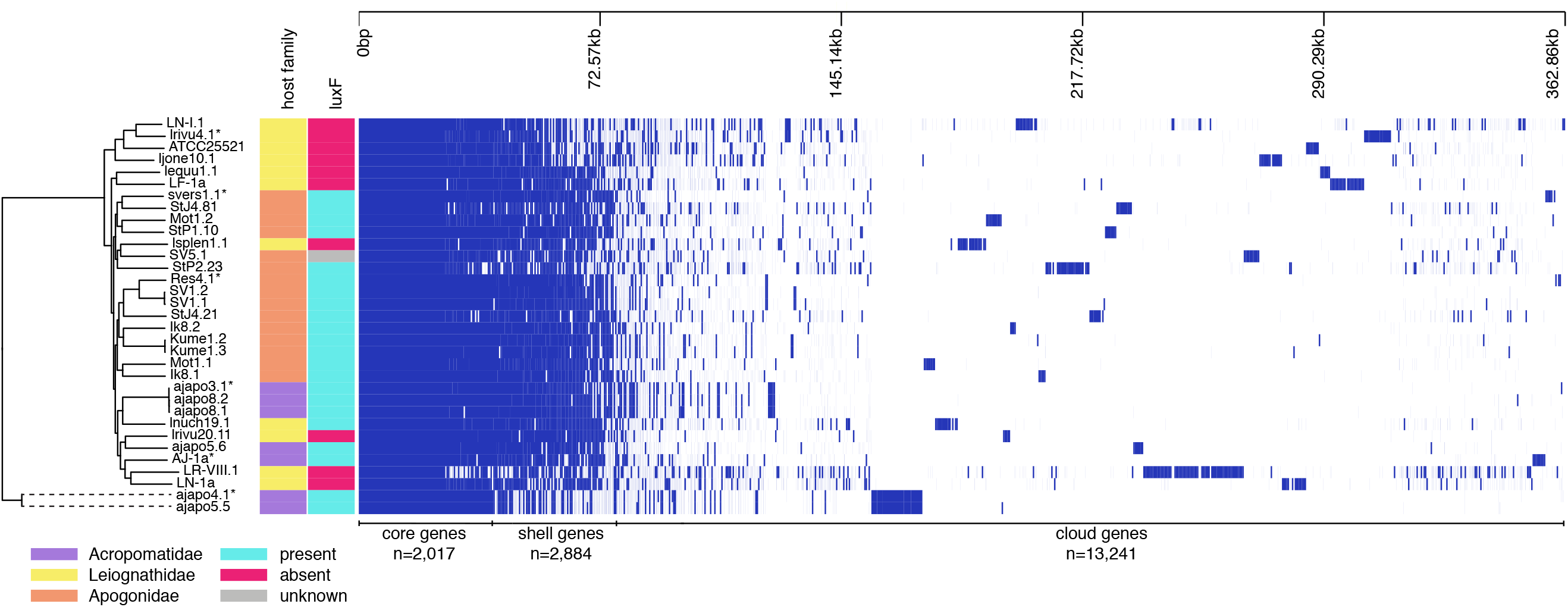
Pangenome analysis of *Photobacterium leiognathi (mandapamensis*) isolated from the light organs of various fish hosts. Phandango plot (Hadfield *et al*. 2018) of gene presence and absence across the core genome the strains where blue indicates the presence of a gene and white indicates its absence. A phylogenetic tree of the strains is also shown as well as each strain’s corresponding host family of origin and the presence or absence of *luxF* in the genome as indicated by the different colors to the right of the phylogeny (see legend for details).

### Phylogenetic Analysis

A phylogenetic analysis based on an alignment of 520 core genes identified across all strains, including the four *P. kishitanii* strains and NCBI reference strains representative of additional *Photobacterium* species, indicates strong support for *P. leiognathi/mandapamensis* clade (Fig. 6a). This analysis also supports the divergence of strains ajapo4.1 and ajapo5.1 from this clade. An additional analysis of these two strains and the *P. leiognathi* and *P. mandapamensis* strains based on an alignment of 2,017 core genes detected in the pangenome analysis indicated 4 distinct clades among the *P. leiognathi* and *P. mandapamensis* strains in this study. Two of the clades are comprised entirely of strains originating from the host fish, *Siphamia tubifer*, with the exception of a single strain, lsplen1.1, which originated from the Leiognathid host, *Eubleekeria splendens*. A third clade is comprised of strains that originated from the light organs of both Leiognathid and Acropomatid fishes, and a fourth clade, basal to the other three, contain only strains originating from Leiognathid hosts, including the *P. leiognathi* type strain ATCC25521 (Fig. 6b). This clade also contains *P. leiognathi* strain lrivu4.1 (GCA_000509205.1), whereas the other three clades contain strains previously identified as *P. mandapamensis*. This analysis also placed ajapo5.5 and ajapo4.1 (GCA_003026025.1) as divergent from the *P. leiognathi* and *P. mandapamensis* strains with high confidence (100/100) (Fig. 5b).

**Figure 6.**
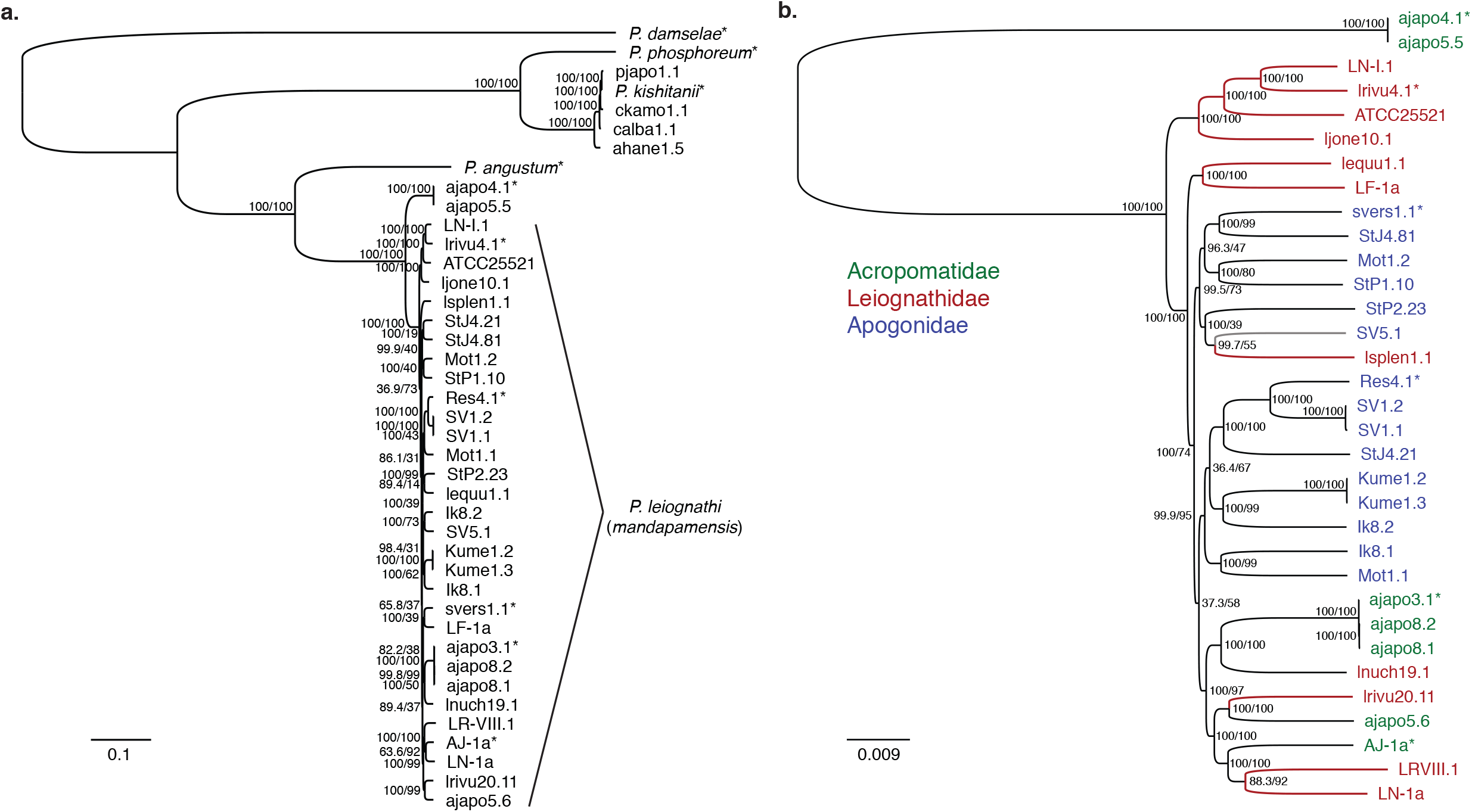
Phylogenetic analysis of *Photobacterium* species isolated from the light organs of various fish hosts. Midpoint rooted trees are shown for an analysis of a) all strains sequenced in this study based on an alignment of 520 core genes with 150 bootstrap replicates and b) only the *P. leiognathi (mandapamensis*) strains based on an alignment of 2,017 core genes with 300 bootstrap replicates. Tip labels are colored according to which family of host fish the strain originated from and the red branches indicates the absence of *luxF* in the genome. Reference strains included in the analyses are indicated by an *. Scale bars show the inferred number of nucleotide substitutions per site. Both trees were constructed using the GTR+F+I+G4 model in IQtree. Values listed on branches indicate bootstrap/SH-aLRT support.

### Lux operon

A comparison of the of the *lux-rib* operon of the different strains revealed a pattern that corresponds with the fish host family from which the bacteria originated. Bacteria isolated from *Acropoma japonicum* (‘ajapo’ strains) and *Siphamia tubifer* hosts all contain the *luxF* gene (Fig. 7). One strain, SV5.1, which was isolated from the light organ of *S. tubifer*, had an incomplete assembly of the *lux* genes and is thus, excluded from this analysis. In contrast, all strains that were isolated from the light organs of Leiognathid fishes, with the exception of one strain, lnuch19.1, did not contain the *luxF* gene, as well as most of the *lumP* gene (Fig. 7). The four *P. kishitanii* isolates all contained *luxF* but lacked both the *lumP* and *lumQ* genes.

**Figure 7.**
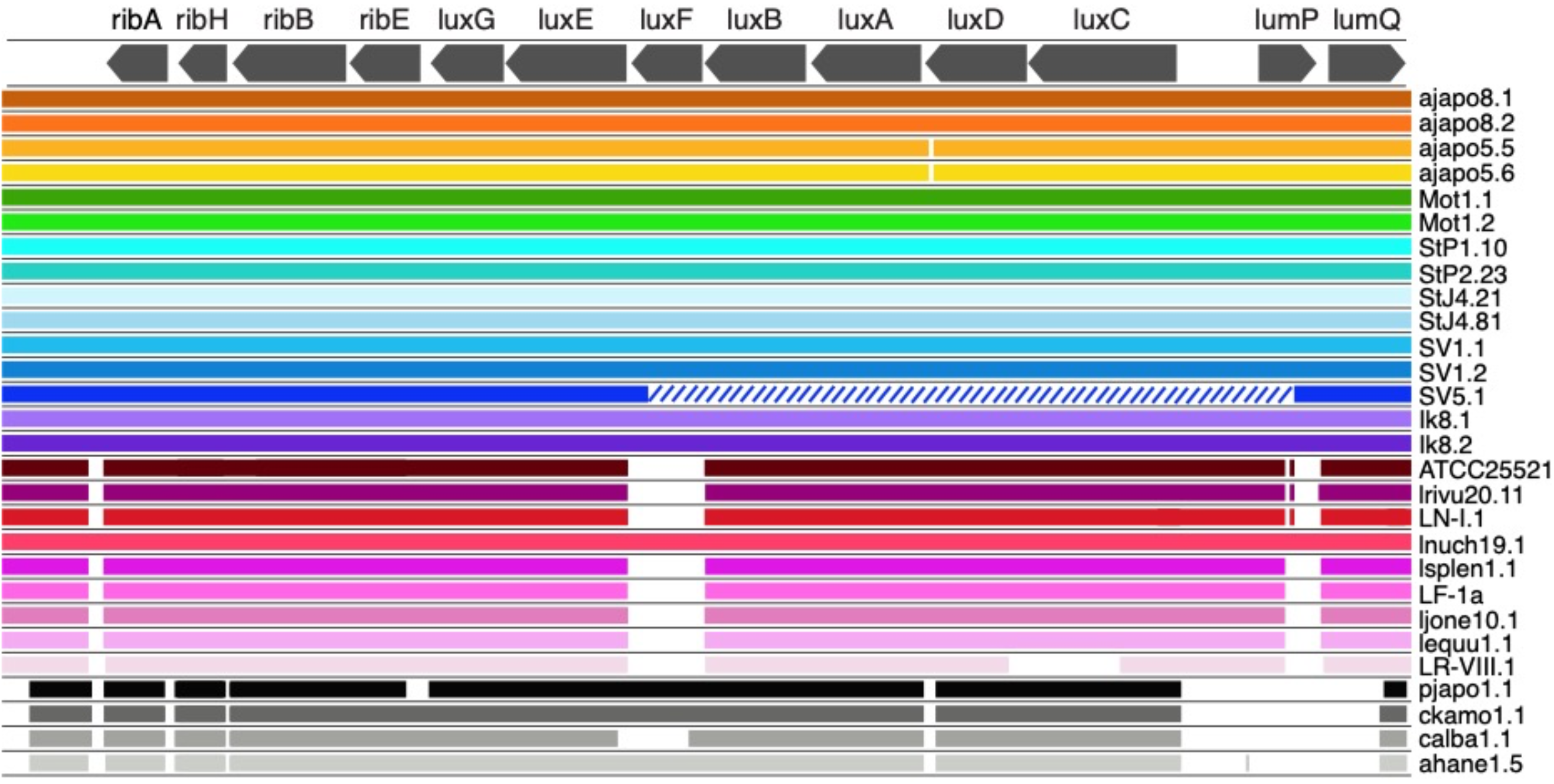
Alignments of the *lux-rib* operon of the *Photobacterium* sp. isolates sequenced in this study. *Photobacterium ‘mandapamensis’* strain svers1.1 (Urbanczyk *et al*. 2011) was used as the reference for BLAST comparisons of the other isolates using an e-value cutoff score of 0.1. The isolate names are listed to the right of their respective colored band. Hash marks indicate an incomplete genome assembly. Figure produced with Proksee (Grant *et al*. 2022).

### Average Nucleotide Identity

The pairwise average nucleotide identity (ANI) analysis across all strains showed a clear distinction between the *P. kishitanii* strains and all others, with an average ANI of 80.6% between the two groups (Fig. 8). The pairwise comparisons among all non-*P. kishitanii* strains resulted in ANIs greater than 95%, with the exception of the two divergent strains mentioned above in the phylogenetic analysis, ajapo5.5 and ajapo4.1. The average ANI between these two strains and the *P. leiognathi* and *P. mandapamensis* strains is 92.9%, below the 95% threshold for bacterial species delimitation (Fig. 8). The average pairwise ANI between the *P. leiognathi* strains and *P. mandapamensis* strains was 96.5% versus 97.3% when the *P. leiognathi* strains were compared to each other.

**Figure 8.**
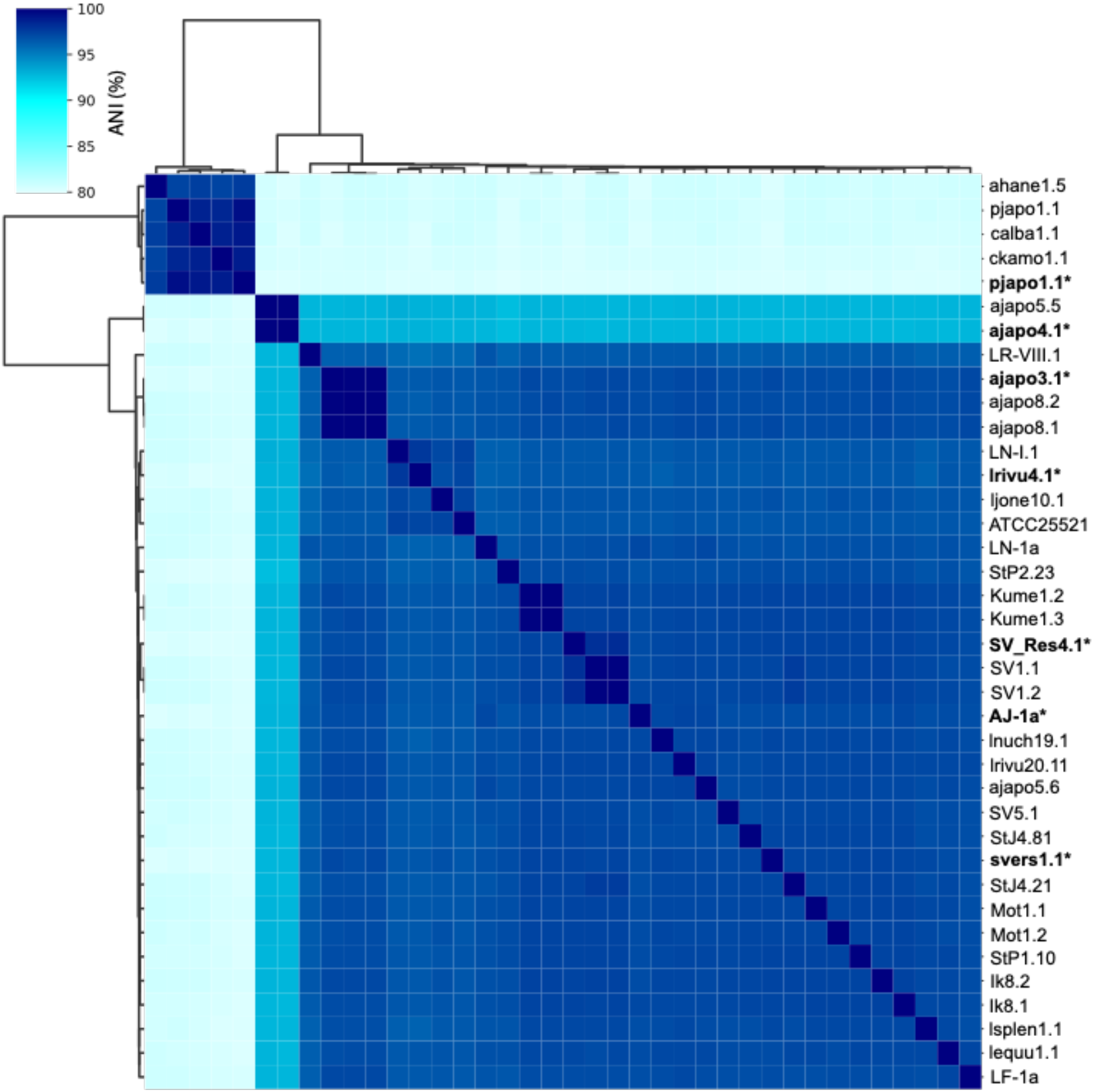
Clustered heatmap depicting the pairwise average nucleotide identities (ANI) across the complete genomes of the *Photobacterium* strains sequenced in this study. Reference strains included in the analysis are in bold and indicated with an *.

### Plasmids

Of the plasmid sequences identified across all strains, 24 were fully circularized. The average length of these circularized sequences ranged from 7,304 to 100,280 bp with a mean of 39,194 bp. Most strains had only one identifiable plasmid, however three circularized plasmids were identified in strains lsplen1.1 and LN-I.1. Four additional strains contained two plasmids. The majority of genes identified across all plasmid sequences were of unknown function, but there were 163 total genes that were assigned function, 30 of which encoded transposases, and 52 that were shared by at least two strains (Fig. 9). The remaining 111 genes were uniquely found in only a single strain (Table S3). Comparing genes across the plasmid sequences from different strains revealed some similarities between plasmids originating from the same host species and location, such as ajapo8.1/ajapo8.2 and Ik8.1/Ik8.2. There were also several genes present in plasmids across all strains, including *bin3, dns*, repA, and *tnpR* (Fig. 9). With respect to transposases, the IS6 family transposase ISPpr9 was the most common in plasmids across strains.

**Figure 9.**
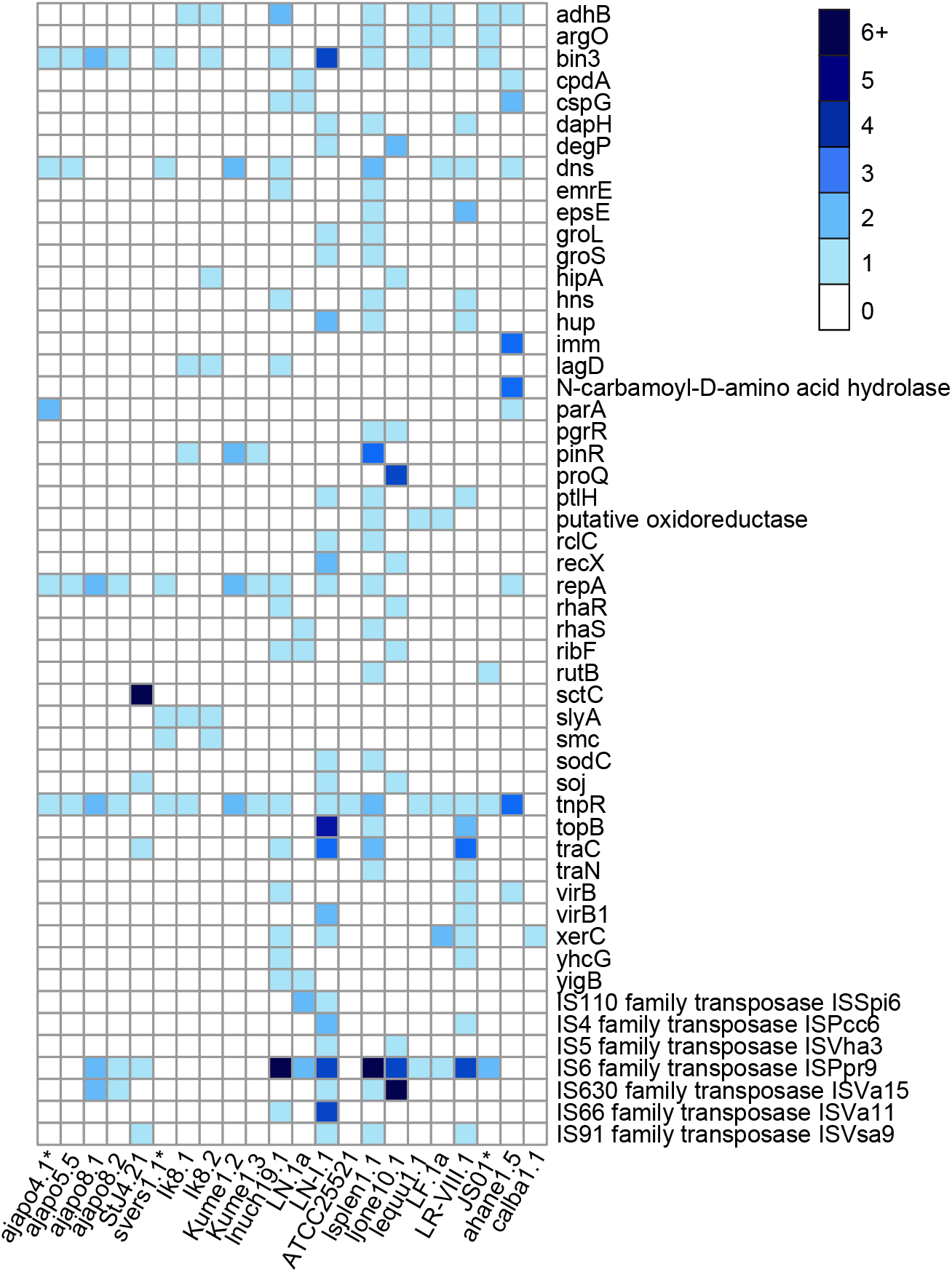
Summary of plasmid gene content of the *Photobacterium* sp. strains sequenced in this study. Genes listed are present in at least two strains and their copy number is indicated by the corresponding legend color. A complete list of genes present in all plasmid sequences identified are presented in Table S3. Reference strains included in the analysis are indicated with a *.

## Discussion

Using ONT sequencing, we were able to assembly highly contiguous genomes of 31 *Photobacterium* species originating from the light organs of 12 species of fish representing 6 unique families in 4 orders. The majority of the assemblies were at the chromosome-level, comprised of one large contig greater than 3 Mbp and one smaller contig approximately 1.5 Mbp. Several strains had additional plasmid sequences ranging in size from approximately 2,000-100,000 bp. These values are consistent with what has been reported for the genomes of other *Photobacterium* species, which are typically comprised of one large and small chromosome between 3.13 to 4.09 Mbp and 1.05 to 2.24 Mbp, respectively, as well as numerous small plasmids (Dunlap *et al*. 2004; Okada *et al*. 2005, Vezzi *et al*. 2005; Kim *et al*. 2008, Urbanczyk *et al*. 2011a). The study represents the largest collection of *Photobacterium* genomes sequenced to date, and more than doubles the number of *P. leiognathi* and *P. mandapamensis* genomes available from NCBI, enabling a more robust analysis of these two groups of closely related lineages.

There are many programs currently available for the assembly of bacterial genomes from ONT sequences and previous studies have compared some of the different assembly approaches (e.g. Goldstein *et al*. 2019, Lee *et al*. 2021, Murigneux *et al*. 2021, Zhang *et al*. 2021), there remains no clear consensus pipeline. Comparing several combinations of various assembly tools used in this study, we chose a pipeline that implemented the Flye assembler (Kolmogorov *et al*. 2019) followed by Circlator (Hunt *et al*. 2015), to orient and circularize the sequences, and both Medak and Homopolish (Huang *et al*. 2021) for polishing. After polishing we ran Ragout (Kolmogorov *et al*. 2014) or RagTag (Algone *et al*. 2021) for scaffolding with varying results depending on the strain. Overall, this pipeline resulted in the assembly of highly contiguous, near-complete *Photobacterium* genomes, even from samples with less than 10x sequence coverage depth. Like other studies (Lee *et al*. 2021, Zhang *et al*. 2021), we found that using Homopolish as a secondary polishing step dramatically improved BUSCO completeness scores, particularly for strains with poor coverage depth. We also saw a reduction in the number of contigs for many assemblies after running Circlator and, most notably, after scaffolding. Not surprisingly, incorporating available short reads to the assembly also increased BUSCO completeness dramatically, although these hybrid assemblies resulted in a larger number of contigs than the long read-only assembly, which were reduced significantly by scaffolding and discarding short contigs (<1,000 bp).

This study confirms that *P. leiognathi* and *P. mandapamensis* are phylogenetically and ecologically closely related and are at the cut-off used to delimit bacterial species, providing a unique opportunity to analyze the early stages of bacterial speciation (Urbanczyk *et al*. 2013). Overall, the differences we see between the *P. leiognathi* and *P. mandapamensis* genomes are minimal, although the *P. leiognathi* strains are approximately 6% larger. The core genome of *P. leiognathi* was also larger than that of *P. mandapamensis*, but this could also be a result of the fact that fewer *P. leiognathi* strains were sequenced in this study. However, it has been suggested that *P. leiognathi* has a more plastic genome and acquires genes horizontally more frequently than *P. mandapamensis* (Urbanczyk *et al*. 2013). In fact, horizontal gene transfer has been shown from more distantly related bacteria to *Photobacterium* species, contributing to the pangenome (Urbanczyk *et al*. 2011). The role of plasmids in the genome evolution of *Photobacterium* has not been thoroughly examined, however, our study confirms previous findings that *Photobacterium* plasmids typically lack essential genes (Campanaro *et al*. 2005). Thus, the role of plasmids in contributing to genetic novelties in *Photobacterium* is likely minimal overall.

Based on the average nucleotide identities of approximately 93% between strains ajapo4.1 and ajapo5.1 and the *P. leiognathi* and *P. mandapamensis* strains, as well as their divergence from the *P. leiognathi/mandapamensis* clade, we suggest that strains ajapo5.5 and ajapo4.1 be considered a new species of *Photobacterium*. We propose the name ‘Candidatus’ *Photobacterium acropomis* to reflect their host of origin, *Acropoma japonicus*. These two strains possess 765 unique genes not present in the other *Photobacterium* genomes. Among these genes are those relating to macrolide export and the production of urease, which catalyzes the hydrolysis of urea. While it remains unclear what the role urease might play for these bacteria, the presence of these genes provides them with the potential to break down urea.

While housekeeping and *lux* gene phylogenies are generally congruent for *Photobacterium* (Urbanczyk *et al*., 2008, 2011), the presence or absence of *luxF* does not correlate with the phylogenic relationships inferred from the core genome assemblies of the *P. leiognathi* and *P. mandapamensis* strains in this study. In fact, several strains inferred to be *P. mandapamensis* based on their phylogenetic positioning are missing *luxF*, suggesting its presence or absence may not be a defining feature between *P. leiognathi* and *P. mandapamensis* (Ast & Dunlap 2004). Interestingly, there is strong congruence between the presence of *luxF* and host family of origin. All strains originating from Apogonid (*Siphamia*) and Acropomatid (*Acropoma*) hosts contained both the *luxF* and *lumP* genes, whereas all strains from Leiognathid hosts were missing both genes, with the exception of one strain, which was most closely related to strains isolated from an *Acropoma* host. While *luxF* is not required for light production, its presence in the *lux* operon appears to increase light emission (Brodl *et al*. 2022), which may be critical for the bioluminescent symbiosis with *Siphamia* and *Acropoma* hosts, and perhaps, is a key genetic feature for these hosts to recognize potential symbionts from the environmental pool of bacteria.

Despite the congruence of host range and the presence of the *luxF* gene, there remains no evidence of the bioluminescent symbiosis having played a role in the divergence of *P. leiognathi* and *P. mandapamensis*. Furthermore, strains from both lineages can be co-symbionts of the same light organ (Kaeding *et al*. 2007). Leiognathid fishes appear to associate with a wide range of both *P. leiognathi* and *P. mandapamensis* strains, as do Acrompomatids, which also associate with ‘Candidatus’ *Photobacterium acropomis* described here. In contrast, Apogonid hosts in the genus *Siphamia* associate with a much narrower range of only *P. mandapamensis* strains (Kaeding *et al*. 2007, Gould *et al*. 2021, this study). It remains unclear why this degree of specificity exists for the bioluminescent symbiosis with *Siphamia* hosts and not for the other hosts examined, but it could be due the host’s distinct behavioral ecology as a cryptic reef fish (Gould *et al*. 2014). Furthermore, there are likely mechanisms in place for the *Siphamia* hosts to identify *P. mandapamensis* from the diverse pool of bacteria in the environment, which could include the presence of *luxF* and other genomic features. As long read sequencing technologies like ONT become more accurate and accessible and new programs are developed to improve genome assembly pipelines, there will be an increasing number of bacterial genomes available to explore the genetic signatures of host-microbe interactions.

## Acknowledgements

We’d like to thank Paul Dunlap for the initial sampling, isolation, and cultivation of many of the strains sequenced in this study. We would also like to thank Athena Lam, Director of the Center for Comparative Genomics at the California Academy of Sciences, for her assistance with Nanopore sequencing.

## Data Accessibility

All genome assemblies will be made publicly available on NCBI and the corresponding scripts used for data analysis will be available on the authors’ Github pages.

## References

Al Ali, B., Garel, M., Cuny, P., Miquel, J. C., Toubal, T., Robert, A., & Tamburini, C. (2010). Luminous bacteria in the deep-sea waters near the ANTARES underwater neutrino telescope (Mediterranean Sea). Chemistry and Ecology, 26(1), 57–72.

Alonge, M., Lebeigle, L., Kirsche, M., Aganezov, S., Wang, X., Lippman, Z. B., … & Soyk, S. (2021). Automated assembly scaffolding elevates a new tomato system for high-throughput genome editing. BioRxiv.

Altschul, S.F., Gish, W., Miller, W., Myers, E.W. & Lipman, D.J. (1990) “Basic local alignment search tool.” J. Mol. Biol. 215: 403–410.

Ast J.C., & Dunlap, P.V. (2004) Phylogenetic analysis of the lux operon distinguishes two evolutionarily distinct clades of Photobacterium leiognathi. Arch Microbiol 181: 352–361.

Ast, J. C., and Dunlap, P. V. (2005). Phylogenetic resolution and habitat specificity of members of the Photobacterium phosphoreum species group. Environmental Microbiology, 7(10), 1641–1654

Ast J.C., Urbanczyk, H., & Dunlap, P.V. (2007) Natural merodiploidy of the lux-rib operon of Photobacterium leiognathi from coastal waters of Honshu, Japan. J Bacteriol 189: 6148–6158.

Ast, J. C., Cleenwerck, I., Engelbeen, K., Urbanczyk, H., Thompson, F. L., De Vos, P., & Dunlap, P. V. (2007). Photobacterium kishitanii sp. nov., a luminous marine bacterium symbiotic with deep-sea fishes. International Journal of Systematic and Evolutionary Microbiology, 57(9), 2073–2078.

Brodl, E., Csamay, A., Horn, C., Niederhauser, J., Weber, H., & Macheroux, P. (2020). The impact of LuxF on light intensity in bacterial bioluminescence. Journal of Photochemistry and Photobiology B: Biology, 207, 111881.

Chen, S., Zhou, Y., Chen, Y., & Gu, J. (2018). fastp: an ultra-fast all-in-one FASTQ preprocessor. Bioinformatics, 34(17), i884–i890.

Dunlap, P. V., Jiemjit, A., Ast, J. C., Pearce, M. M., Marques, R. R., & Lavilla-Pitogo, C. R. (2004). Genomic polymorphism in symbiotic populations of Photobacterium leiognathi. Environmental Microbiology, 6(2), 145–158.

Dunlap, P. V., & Ast, J. C. (2005). Genomic and phylogenetic characterization of luminous bacteria symbiotic with the deep-sea fish Chlorophthalmus albatrossis (Aulopiformes: Chlorophthalmidae). Applied and environmental microbiology, 71(2), 930–939.

Fukasawa, S., Suda, T., & Kubota, S. (1988). Identification of luminous bacteria isolated from the light organ of the fish, Acropoma japonicum. Agricultural and biological chemistry, 52(1), 285–286.

Goldstein, S., Beka, L., Graf, J., & Klassen, J. L. (2019). Evaluation of strategies for the assembly of diverse bacterial genomes using MinION long-read sequencing. BMC genomics, 20(1), 1–17.

Gould, A. L., Harii, S., & Dunlap, P. V. (2014). Host preference, site fidelity, and homing behavior of the symbiotically luminous cardinalfish, Siphamia tubifer (Perciformes: Apogonidae). Marine biology, 161(12), 2897–2907.

Gould, A. L., Fritts-Penniman, A., & Gaisiner, A. (2021). Museum genomics illuminate the high specificity of a bioluminescent symbiosis for a genus of reef fish. Frontiers in ecology and evolution, 9, 630207.

Gould, A.L., Donohoo, S., & Neff, E. (*in prep*) Distinct strain-level symbiont communities between individuals and populations of a bioluminescent fish host

Grant, J.R., Enns, E., Marinier, E., Saha-Mandal, A., Chen, C-Y., Graham, M., Van Domselaar, G., & Stothard, P. (2022) A web server for assembling, annotating and visualizing bacterial genomes. Poster, BioNet 2022 Conference

Gurevich, A., Saveliev, V., Vyahhi, N., & Tesler, G. (2013). QUAST: quality assessment tool for genome assemblies. Bioinformatics, 29(8), 1072–1075.

Hadfield, J., Croucher, N. J., Goater, R. J., Abudahab, K., Aanensen, D. M., & Harris, S. R. (2018). Phandango: an interactive viewer for bacterial population genomics. Bioinformatics, 34(2), 292–293.

Huang, Y. T., Liu, P. Y., & Shih, P. W. (2021). Homopolish: a method for the removal of systematic errors in nanopore sequencing by homologous polishing. Genome biology, 22(1), 1–17.

Hunt, M., Silva, N. D., Otto, T. D., Parkhill, J., Keane, J. A., & Harris, S. R. (2015). Circlator: automated circularization of genome assemblies using long sequencing reads. Genome biology, 16(1), 1–10.

Jain, C., Rodriguez-R, L. M., Phillippy, A. M., Konstantinidis, K. T., & Aluru, S. (2018). High throughput ANI analysis of 90K prokaryotic genomes reveals clear species boundaries. Nature communications, 9(1), 1–8.

Kaeding, A. J., Ast, J. C., Pearce, M. M., Urbanczyk, H., Kimura, S., Endo, H., … & Dunlap, P. V. (2007). Phylogenetic diversity and cosymbiosis in the bioluminescent symbioses of “Photobacterium mandapamensis”. Applied and Environmental Microbiology, 73(10), 3173–3182.

Kolmogorov, M., Raney, B., Paten, B., & Pham, S. (2014). Ragout—a reference-assisted assembly tool for bacterial genomes. Bioinformatics, 30(12), i302–i309.

Kolmogorov, M., Yuan, J., Lin, Y., & Pevzner, P. A. (2019). Assembly of long, error-prone reads using repeat graphs. Nature biotechnology, 37(5), 540–546.

Lee, J. Y., Kong, M., Oh, J., Lim, J., Chung, S. H., Kim, J. M., … & Kwak, W. (2021). Comparative evaluation of Nanopore polishing tools for microbial genome assembly and polishing strategies for downstream analysis. Scientific Reports, 11(1), 1–11.

Nguyen, L. T., Schmidt, H. A., Von Haeseler, A., & Minh, B. Q. (2015). IQ-TREE: a fast and effective stochastic algorithm for estimating maximum-likelihood phylogenies. Molecular biology and evolution, 32(1), 268–274.

Okada, K., Iida, T., Kita-Tsukamoto, K., & Honda, T. (2005). Vibrios commonly possess two chromosomes. Journal of bacteriology, 187(2), 752–757.

Page, A. J., Cummins, C. A., Hunt, M., Wong, V. K., Reuter, S., Holden, M. T., … & Parkhill, J. (2015). Roary: rapid large-scale prokaryote pan genome analysis. Bioinformatics, 31(22), 3691–3693.

Seemann, T. (2014). Prokka: rapid prokaryotic genome annotation. Bioinformatics, 30(14), 2068–2069.

Seppey, M., Manni, M., & Zdobnov, E. M. (2019). BUSCO: assessing genome assembly and annotation completeness. In Gene prediction (pp. 227–245). Humana, New York, NY.

Shimoyama, Y. (2022) ANIclustermap: A tool for drawing ANI clustermap between all-vs-all microbial genomes (https://github.com/moshi4/ANIclustermap)

Soh, J. Y., Russell, C. W., Fenlon, S. N., & Chen, S. L. (2018). Complete genome sequence of Photobacterium leiognathi strain JS01. Genome Announcements, 6(1), e01396–17.

Urbanczyk, H., Ast, J. C., Kaeding, A. J., Oliver, J. D., & Dunlap, P. V. (2008). Phylogenetic analysis of the incidence of lux gene horizontal transfer in Vibrionaceae. Journal of bacteriology, 190(10), 3494–3504.

Urbanczyk, H., Ast, J. C., & Dunlap, P. V. (2011a). Phylogeny, genomics, and symbiosis of Photobacterium. FEMS microbiology reviews, 35(2), 324–342.

Urbanczyk, H., Ogura, Y., Hendry, T. A., Gould, A. L., Kiwaki, N., Atkinson, J. T., … & Dunlap, P. V. (2011b). Genome sequence of Photobacterium mandapamensis strain svers.1.1, the bioluminescent symbiont of the cardinal fish Siphamia versicolor.

Urbanczyk, H., Urbanczyk, Y., Hayashi, T., & Ogura, Y. (2013). Diversification of two lineages of symbiotic Photobacterium. PloS one, 8(12), e82917.

Wada, M., Kamiya, A., Uchiyama, N., Yoshizawa, S., Kita-Tsukamoto, K., Ikejima, K., … & Kogure, K. (2006). Lux A gene of light organ symbionts of the bioluminescent fish Acropoma japonicum (Acropomatidae) and Siphamia versicolor (Apogonidae) forms a lineage closely related to that of Photobacterium leiognathi ssp. Mandapamensis. FEMS microbiology letters, 260(2), 186–192.

Wick, R. R., Judd, L. M., Gorrie, C. L., & Holt, K. E. (2017). Unicycler: resolving bacterial genome assemblies from short and long sequencing reads. PLoS computational biology, 13(6), e1005595.

Zhang, P., Jiang, D., Wang, Y., Yao, X., Luo, Y., & Yang, Z. (2021). Comparison of de novo assembly strategies for bacterial genomes. International Journal of Molecular Sciences, 22(14), 7668.

