## Supplemental Tables for "Highly contiguous genome assemblies of *Photobacterium* strains isolated from fish light organs using nanopore sequencing technology"

**Summary**

This document contains the supplemental tables (S1-3) for the manuscript: “Highly contiguous genome assemblies of *Photobacterium* strains isolated from fish light organs using nanopore sequencing technology” by AL Gould and JB Henderson.

**Table S1**. Summary of draft assemblies before and after scaffolding. Listed are the number of contigs, the total number of bp in the assembly, the largest contig size (bp), the GC content, N50 and L50 values, the number of coding sequences (CDS), the number of rRNAs and tRNAs and the BUSCO scores: complete [single, duplicate], fragmented, and missing.

| **Strain** | **Assembly** | **Ctgs** | **Total bp** | **GC%** | **N50** | **L50** | **Ns** | **CDS** | **rRNA** | **tRNA** | **Complete [S, D]** | **Fragment** | **Missing** |
| --- | --- | --- | --- | --- | --- | --- | --- | --- | --- | --- | --- | --- | --- |
| pjapo1.1 | pre-scaffold | 38 | 4389259 | 39.36 | 286117 | 5 | 0 | 5960 | 25 | 196 | 66.1% [64.8%, 1.3%] | 12.10% | 21.80% |
|  | ragtag scaffold | 4 | 4392659 | 39.36 | 3165177 | 1 | 77 | 5932 | 25 | 1 | 66.1% [64.8%, 1.3%] | 12.00% | 21.90% |
|  | **ragout scaffold** | **4** | **5000498** | **39.36** | **3339696** | **1** | **12224** | **6596** | **25** | **196** | **66.1% [64.8%, 1.3%]** | **12.00%** | **21.90%** |
| ckamo1.1 | pre-scaffold | 27 | 4913886 | 38.96 | 481211 | 4 | 0 | 6183 | 8 | 173 | 74.0% [73.2%, 0.8%] | 9.50% | 16.50% |
|  | ragtag scaffold | 4 | 4916186 | 38.96 | 3232254 | 1 | 47 | 6027 | 8 | 173 | 74.0% [73.2%, 0.8%] | 9.50% | 16.50% |
|  | **ragout scaffold** | **4** | **5039339** | **38.96** | **3317370** | **1** | **2489** | **6136** | **8** | **173** | **74.0% [73.2%, 0.8%]** | **9.50%** | **16.50%** |
| ahane1.5 | pre-scaffold | 8 | 5094422 | 39.02 | 2927850 | 1 | 0 | 4694 | 53 | 207 | 95.2% [90.5%, 4.7%] | 1.60% | 3.20% |
|  | ragtag scaffold | 5 | 5094722 | 39.02 | 3347557 | 1 | 6 | 4676 | 53 | 207 | 95.2% [90.5%, 4.7%] | 1.60% | 3.20% |
|  | **ragout scaffold** | **5** | **5100844** | **39.02** | **3353657** | **1** | **126** | **4680** | **53** | **207** | **95.2% [90.5%, 4.7%]** | **1.60%** | **3.20%** |
| calba1.1 | pre-scaffold | 16 | 5159790 | 38.96 | 840622 | 3 | 0 | 4895 | 46 | 206 | 95.0% [94.7%, 0.3%] | 1.50% | 3.50% |
|  | ragtag scaffold | 7 | 5160690 | 38.96 | 3249927 | 1 | 17 | 4880 | 46 | 206 | 95.0% [94.7%, 0.3%] | 1.50% | 3.50% |
|  | **ragout scaffold** | **7** | **5200973** | **38.96** | **3263220** | **1** | **792** | **4892** | **46** | **206** | **95.0% [94.7%, 0.3%]** | **1.50%** | **3.50%** |
| Mot1.1 | pre-scaffold | 8 | 4676157 | 41.25 | 1938146 | 2 | 0 | 4066 | 62 | 208 | 98.5% [98.1%, 0.4%] | 0.20% | 1.30% |
|  | **ragtag scaffold** | **2** | **4676757** | **41.25** | **3973505** | **1** | **13** | **4069** | **62** | **208** | **98.5% [98.1%, 0.4%]** | **0.20%** | **1.30%** |
|  | ragout scaffold | 7 | 4676179 | 41.25 | 3973427 | 1 | 0 | 4069 | 62 | 208 | 98.5% [98.1%, 0.4%] | 0.20% | 1.30% |
| lnuch19.1 | pre-scaffold | 8 | 5229188 | 41.18 | 1534541 | 2 | 0 | 4566 | 57 | 197 | 98.8% [98.2%, 0.6%] | 0.50% | 0.70% |
|  | ragtag scaffold | - | - | - | - | - | - | - | - | - | - | - | - |
|  | **ragout scaffold** | **5** | **5237683** | **41.18** | **3439577** | **1** | **162** | **4618** | **57** | **197** | **98.7% [98.1%, 0.6%]** | **0.50%** | **0.80%** |
| ajapo8.2 | pre-scaffold | 3 | 4886267 | 41.15 | 3261968 | 1 | 0 | 4294 | 53 | 194 | 98.8% [98.3%, 0.5%] | 0.30% | 0.90% |
|  | ragtag scaffold | 3 | 4886267 | 41.15 | 3261968 | 1 | 0 | 4240 | 53 | 194 | 98.8% [98.3%, 0.5%] | 0.40% | 1.10% |
|  | **ragout scaffold** | **3** | **4886267** | **41.15** | **3261968** | **1** | **0** | **4239** | **53** | **194** | **98.8% [98.3%, 0.5%]** | **0.30%** | **0.90%** |
| lsplen1.1 | pre-scaffold | 14 | 5249460 | 41.01 | 1177766 | 2 | 0 | 4818 | 37 | 174 | 98.3% [97.8%, 0.5%] | 0.60% | 1.10% |
|  | ragtag scaffold | 7 | 5250160 | 41.01 | 3241604 | 1 | 13 | 4756 | 37 | 174 | 98.3% [97.8%, 0.5%] | 0.60% | 1.60% |
|  | **ragout scaffold** | **7** | **5292468** | **41.01** | **3284001** | **1** | **813** | **4768** | **37** | **174** | **98.3% [97.8%, 0.5%]** | **0.60%** | **1.10%** |
| Mot1.2 | pre-scaffold | 8 | 4820778 | 41.2 | 794793 | 2 | 0 | 4289 | 62 | 206 | 97.3% [97.0%, 0.3%] | 1.10% | 1.60% |
|  | ragtag scaffold | 2 | 4821378 | 41.2 | 3197395 | 1 | 12 | 4317 | 62 | 206 | 97.3% [97.0%, 0.3%] | 1.10% | 1.60% |
|  | **ragout scaffold** | **2** | **4947312** | **41.2** | **3234211** | **1** | **2558** | **4336** | **62** | **206** | **97.3% [97.0%, 0.3%]** | **1.10%** | **1.60%** |
| LN-I.1 | pre-scaffold | 26 | 5489529 | 41.53 | 715204 | 3 | 0 | 4995 | 62 | 200 | 98.0% [97.4%, 0.6%] | 0.60% | 1.40% |
|  | **ragtag scaffold** | **15** | **5490629** | **41.53** | **3405483** | **1** | **20** | **4994** | **62** | **200** | **98.0% [97.4%, 0.6%]** | **0.60%** | **1.40%** |
|  | ragout scaffold | 16 | 5589437 | 41.53 | 3496794 | 1 | 1787 | 5017 | 62 | 200 | 98.0% [97.4%, 0.6%] | 0.60% | 1.40% |
| Kume1.2 | pre-scaffold | 4 | 4756280 | 41.25 | 3035257 | 1 | 0 | 4208 | 49 | 197 | 98.8% [98.5%, 0.3%] | 0.30% | 0.90% |
|  | ragtag scaffold | 3 | 4756380 | 41.25 | 3186856 | 1 | 2 | 4145 | 49 | 197 | 98.8% [98.5%, 0.3%] | 0.60% | 1.10% |
|  | **ragout scaffold** | **3** | **4835311** | **41.25** | **3186767** | **1** | **1634** | **4156** | **49** | **197** | **98.8% [98.5%, 0.3%]** | **0.30%** | **0.90%** |
| SV1.2 | **pre-scaffold** | **3** | **4718342** | **41.17** | **3184478** | **1** | **0** | **4083** | **51** | **200** | **98.5% [98.1%, 0.4%]** | **0.30%** | **1.20%** |
|  | ragtag scaffold | 2 | 4718442 | 41.17 | 3184478 | 1 | 2 | 4082 | 51 | 200 | 98.5% [98.1%, 0.4%] | 0.30% | 1.20% |
|  | ragout scaffold | 2 | 4718353 | 41.17 | 3184478 | 1 | 0 | 4083 | 51 | 200 | 98.5% [98.1%, 0.4%] | 0.30% | 1.20% |
| Ik8.1 | pre-scaffold | 3 | 4765281 | 41.15 | 3231342 | 1 | 0 | 4127 | 52 | 202 | 98.9% [98.6%, 0.3%] | 0.20% | 0.90% |
|  | ragtag scaffold | 3 | 4765281 | 41.15 | 3231342 | 1 | 0 | 4129 | 47 | 197 | 98.9% [98.6%, 0.3%] | 0.20% | 0.90% |
|  | **ragout scaffold** | **3** | **4765281** | **41.15** | **3231342** | **1** | **0** | **4130** | **47** | **197** | **98.9% [98.6%, 0.3%]** | **0.20%** | **0.90%** |
| Kume1.3 | pre-scaffold | 5 | 4726851 | 41.23 | 2926205 | 1 | 0 | 4177 | 43 | 194 | 98.4% [98.1%, 0.3%] | 0.50% | 1.10% |
|  | ragtag scaffold | 3 | 4727051 | 41.23 | 3172182 | 1 | 4 | 4127 | 43 | 194 | 98.4% [98.1%, 0.3%] | 0.50% | 1.10% |
|  | **ragout scaffold** | **3** | **4818955** | **41.23** | **3185044** | **1** | **1911** | **4137** | **43** | **194** | **98.4% [98.1%, 0.3%]** | **0.50%** | **1.10%** |
| StP2.23 | pre-scaffold | 38 | 4202604 | 41.09 | 209731 | 7 | 0 | 4219 | 27 | 152 | 77.8% [77.7%, 0.1%] | 3.60% | 18.60% |
|  | ragtag scaffold | 6 | 4205804 | 41.09 | 2658946 | 1 | 76 | 4211 | 20 | 144 | 77.8% [77.7%, 0.1%] | 3.60% | 18.60% |
|  | **ragout scaffold** | **4** | **4888897** | **41.09** | **3236904** | **1** | **14038** | **4525** | **20** | **144** | **77.8% [77.7%, 0.1%]** | **3.60%** | **18.60%** |
| StJ4.21 | pre-scaffold | 11 | 4847051 | 41.23 | 929383 | 3 | 0 | 4346 | 63 | 208 | 95.8% [95.5%, 0.3%] | 1.40% | 2.80% |
|  | ragtag scaffold | 4 | 4847751 | 41.23 | 3198579 | 1 | 14 | 4351 | 63 | 208 | 95.8% [95.5%, 0.3%] | 1.40% | 2.80% |
|  | **ragout scaffold** | **4** | **4877430** | **41.23** | **3228436** | **1** | **623** | **4361** | **63** | **208** | **95.8% [95.5%, 0.3%]** | **1.40%** | **2.80%** |
| SV1.1 | pre-scaffold | 8 | 4687119 | 41.13 | 2918729 | 1 | 0 | 4180 | 38 | 190 | 98.1% [97.5%, 0.6%] | 0.60% | 1.30% |
|  | ragtag scaffold | 2 | 4687719 | 41.13 | 3165485 | 1 | 13 | 4116 | 38 | 190 | 98.1% [97.5%, 0.6%] | 0.60% | 1.30% |
|  | **ragout scaffold** | **2** | **4749844** | **41.13** | **3174417** | **1** | **1321** | **4126** | **38** | **190** | **98.1% [97.5%, 0.6%]** | **0.60%** | **1.30%** |
| StP1.10 | pre-scaffold | 12 | 4780150 | 41.15 | 470369 | 3 | 0 | 4189 | 47 | 194 | 98.8% [98.5%, 0.3%] | 0.30% | 0.90% |
|  | ragtag scaffold | 3 | 4781050 | 41.15 | 3175608 | 1 | 19 | 4194 | 47 | 194 | 98.8% [98.5%, 0.3%] | 0.30% | 0.90% |
|  | **ragout scaffold** | **4** | **4853990** | **41.15** | **3193844** | **1** | **1521** | **4208** | **47** | **194** | **98.8% [98.5%, 0.3%]** | **0.30%** | **0.90%** |
| StJ4.81 | pre-scaffold | 42 | 4610923 | 41.33 | 215021 | 6 | 0 | 4134 | 46 | 190 | 94.5% [93.3%, 1.2%] | 1.60% | 3.90% |
|  | ragtag scaffold | 3 | 4614823 | 41.33 | 3008273 | 1 | 85 | 4279 | 46 | 187 | 94.6% [93.4%, 1.2%] | 1.60% | 3.80% |
|  | **ragout scaffold** | **3** | **4967967** | **41.33** | **3223230** | **1** | **7187** | **4378** | **46** | **187** | **94.4% [93.2%, 1.2%]** | **1.60%** | **4.00%** |
| ajapo8.1 | pre-scaffold | 4 | 4878649 | 41.12 | 3245602 | 1 | 0 | 4281 | 44 | 190 | 98.5% [97.9%, 0.6%] | 0.70% | 0.80% |
|  | ragtag scaffold | 4 | 4878649 | 41.12 | 3245602 | 1 | 0 | 4254 | 44 | 192 | 98.5% [97.9%, 0.6%] | 0.70% | 0.80% |
|  | **ragout scaffold** | **4** | **4878649** | **41.12** | **3245602** | **1** | **0** | **4255** | **44** | **192** | **98.5% [97.9%, 0.6%]** | **0.70%** | **0.80%** |
| ajapo5.6 | pre-scaffold | 2 | 4729792 | 41.27 | 3174233 | 1 | 0 | 4104 | 62 | 213 | 98.7% [98.4%, 0.3%] | 0.20% | 1.10% |
|  | ragtag scaffold | 2 | 4729792 | 41.27 | 3174233 | 1 | 0 | 4052 | 62 | 213 | 98.7% [98.4%, 0.3%] | 0.20% | 1.10% |
|  | **ragout scaffold** | **2** | **4729792** | **41.27** | **3174233** | **1** | **0** | **4052** | **62** | **213** | **98.7% [98.4%, 0.3%]** | **0.20%** | **1.10%** |
| LR-VIII.1 | pre-scaffold | 105 | 4467508 | 41.28 | 64254 | 20 | 0 | 4593 | 32 | 147 | 76.6% [76.3%, 0.3%] | 4.40% | 19.00% |
|  | ragtag scaffold | **-** | **-** | **-** | **-** | **-** | **-** | - | - | - | - | - | - |
|  | **ragout scaffold** | **21** | **5791416** | **41.28** | **3699906** | **1** | **22860** | **5602** | **32** | **147** | **76.6% [76.3%, 0.3%]** | **4.20%** | **19.20%** |
| SV5.1 | pre-scaffold | 43 | 4516983 | 41.27 | 190888 | 8 | 0 | 4123 | 35 | 193 | 94.6% [94.3%, 0.3%] | 1.20% | 4.20% |
|  | ragtag scaffold | 2 | 4521083 | 41.27 | 3089627 | 1 | 91 | 4122 | 35 | 193 | 94.7% [94.4%, 0.3%] | 1.20% | 4.10% |
|  | **ragout scaffold** | **3** | **5111110** | **41.27** | **3324477** | **1** | **11624** | **4280** | **35** | **193** | **94.6% [94.3%, 0.3%]** | **1.20%** | **4.20%** |
| ljone10.1 | pre-scaffold | 9 | 5276659 | 41.34 | 3074291 | 1 | 0 | 4551 | 62 | 207 | 98.7% [98.3%, 0.4%] | 0.10% | 1.20% |
|  | ragtag scaffold | - | - | - | - | - | - | - | - | - | - | - | - |
|  | **ragout scaffold** | **8** | **5276714** | **41.34** | **3313411** | **1** | **1** | **4535** | **62** | **207** | **98.7% [98.3%, 0.4%]** | **0.10%** | **1.20%** |
| LN-1a | pre-scaffold | 38 | 5171490 | 41.21 | 359929 | 5 | 0 | 4666 | 44 | 197 | 96.1% [95.8%, 0.3%] | 1.00% | 2.90% |
|  | **ragtag scaffold** | **10** | **5174290** | **41.21** | **3359026** | **1** | **54** | **4760** | **44** | **195** | **96.0% [95.7%, 0.3%]** | **1.10%** | **2.90%** |
|  | ragout scaffold | 12 | 5578355 | 41.21 | 3411149 | 1 | 7294 | 4939 | 44 | 195 | 96.1% [95.8%, 0.3%] | 1.00% | 2.90% |
| ATCC25521 | **pre-scaffold** | **3** | **4750881** | **41.02** | **3269131** | **1** | **0** | **4257** | **47** | **199** | **96.3% [95.9%, 0.4%]** | **1.60%** | **2.10%** |
|  | ragtag scaffold | 3 | 4750881 | 41.02 | 3269131 | 1 | 0 | 4254 | 41 | 194 | 96.3% [95.9%, 0.4%] | 1.70% | 2.10% |
|  | ragout scaffold | 3 | 4750892 | 41.02 | 3269131 | 1 | 0 | 4253 | 41 | 194 | 96.3% [95.9%, 0.4%] | 1.60% | 2.10% |
| lequu1.1 | **pre-scaffold** | **4** | **4825624** | **41.19** | **3204041** | **1** | **0** | **4206** | **62** | **209** | **98.7% [98.3%, 0.4%]** | **0.30%** | **1.00%** |
|  | ragtag scaffold | - | - | - | - | - | - | - | - | - | - | - | - |
|  | ragout scaffold | 4 | 4825624 | 41.19 | 3204041 | 1 | 0 | 4192 | 62 | 209 | 98.7% [98.3%, 0.4%] | 0.30% | 1.00% |
| LF-1a | pre-scaffold | 13 | 5042264 | 40.96 | 915751 | 2 | 0 | 4575 | 36 | 189 | 98.5% [97.9%, 0.6%] | 0.50% | 1.00% |
|  | ragtag scaffold | 6 | 5042964 | 40.96 | 3186469 | 1 | 14 | 4593 | 39 | 192 | 98.5% [97.9%, 0.6%] | 1.80% | 2.70% |
|  | **ragout scaffold** | **5** | **5104659** | **40.96** | **3267821** | **1** | **1222** | **4607** | **39** | **192** | **98.5% [97.9%, 0.6%]** | **0.50%** | **1.00%** |

**Table S2**. List of unique genes found in *Photobacterium* strains ajapo4.1 and ajapo5.5 with available annotations as determined by the pangenome analysis with Roary (Page *et al.* 2015).

| **ID** | **Gene** | **Product** |
| --- | --- | --- |
| group_9693 | aat | Leucyl/phenylalanyl-tRNA--protein transferase |
| group_10271 | aceK | Isocitrate dehydrogenase kinase/phosphatase |
| group_9760 | ackA_2 | Acetate kinase |
| group_17944 | adhB_3 | Alcohol dehydrogenase 2 |
| group_10178 | aes | Acetyl esterase |
| group_10196 | agaS | D-galactosamine-6-phosphate deaminase AgaS |
| group_10020 | ahpD | Alkyl hydroperoxide reductase AhpD |
| group_10014 | ald_2 | Alanine dehydrogenase |
| group_9717 | amiC | Aliphatic amidase expression-regulating protein |
| group_10127 | ART1 | Putative NAD(+)--arginine ADP-ribosyltransferase Vis |
| group_10090 | ASPA | aspartoacylase |
| group_9965 | astB | N-succinylarginine dihydrolase |
| group_9571 | bacC | Dihydroanticapsin 7-dehydrogenase |
| group_10136 | baiA | 3-alpha-hydroxycholanate dehydrogenase (NADP(+)) |
| group_9951 | bepF_1 | Efflux pump periplasmic linker BepF |
| group_9768 | betA | Oxygen-dependent choline dehydrogenase |
| bfr | bfr | Bacterioferritin |
| group_9797 | birA | Bifunctional ligase/repressor BirA |
| group_9813 | bkdB | Lipoamide acyltransferase component of branched-chain alpha-keto acid dehydrogenase complex |
| group_9513 | blc | Outer membrane lipoprotein Blc |
| group_9865 | bmr3 | Multidrug resistance protein 3 |
| group_9541 | btuC | Vitamin B12 import system permease protein BtuC |
| btuD_3 | btuD_3 | Vitamin B12 import ATP-binding protein BtuD |
| group_9628 | btuF_1 | Vitamin B12-binding protein |
| group_10274 | btuF_2 | Vitamin B12-binding protein |
| group_9484 | cadC | Transcriptional activator CadC |
| group_10216 | cheB | Protein-glutamate methylesterase/protein-glutamine glutaminase |
| group_9803 | Chelt | NAD(+)--arginine ADP-ribosyltransferase Chelt |
| group_9863 | cheV_2 | Chemotaxis protein CheV |
| group_9545 | chiA_2 | Chitinase A |
| group_10171 | chiA_3 | Chitinase A |
| group_9847 | cntI_2 | Pseudopaline exporter CntI |
| group_9850 | cobC | Adenosylcobalamin/alpha-ribazole phosphatase |
| group_9849 | cobP | Bifunctional adenosylcobalamin biosynthesis protein CobP |
| group_9851 | cobQ | Cobyric acid synthase |
| group_10241 | cpxP | Periplasmic protein CpxP |
| group_9844 | cqsS | CAI-1 autoinducer sensor kinase/phosphatase CqsS |
| group_9972 | cry1 | Cryptochrome DASH |
| cysA_1 | cysA_1 | Putative thiosulfate sulfurtransferase |
| group_9871 | cysG_1 | Siroheme synthase |
| group_6431 | damX_2 | hypothetical protein |
| group_9747 | dapH_1 | 2,3,4,5-tetrahydropyridine-2,6-dicarboxylate N-acetyltransferase |
| group_9700 | dgcJ | putative diguanylate cyclase DgcJ |
| group_9677 | dinG_2 | 3'-5' exonuclease DinG |
| group_9895 | dinI | DNA damage-inducible protein I |
| group_9602 | dltA | D-alanine--D-alanyl carrier protein ligase |
| group_9563 | dmlR_5 | HTH-type transcriptional regulator DmlR |
| group_9885 | dmlR_8 | HTH-type transcriptional regulator DmlR |
| group_10235 | eamB | Cysteine/O-acetylserine efflux protein |
| group_9565 | emrE_1 | Multidrug transporter EmrE |
| group_10242 | envC | Murein hydrolase activator EnvC |
| group_9593 | epsC | Type II secretion system protein C |
| epsL_2 | epsL_2 | putative sugar transferase EpsL |
| group_10145 | eptA_3 | Phosphoethanolamine transferase EptA |
| exsA | exsA | Exoenzyme S synthesis regulatory protein ExsA |
| group_10179 | fabV_2 | Trans-2-enoyl-CoA reductase [NADH] |
| group_10040 | fan1 | Fanconi-associated nuclease 1 |
| group_9756 | fepE | Ferric enterobactin transport protein FepE |
| group_10159 | fetA | putative iron export ATP-binding protein FetA |
| group_9664 | fic | putative protein adenylyltransferase Fic |
| group_10240 | ftsN | Cell division protein FtsN |
| group_10015 | gcvA_2 | Glycine cleavage system transcriptional activator |
| group_10135 | gcvA_2 | Glycine cleavage system transcriptional activator |
| group_9622 | gcvH_2 | Glycine cleavage system H protein |
| group_9807 | gcvH_2 | Glycine cleavage system H protein |
| group_1539 | gcvP | Glycine dehydrogenase (decarboxylating) |
| group_9632 | ghrA | Glyoxylate/hydroxypyruvate reductase A |
| group_9588 | glpE_1 | Thiosulfate sulfurtransferase GlpE |
| group_9938 | gltC_1 | HTH-type transcriptional regulator GltC |
| group_9993 | gph_2 | Phosphoglycolate phosphatase |
| group_9852 | gpmA | 2,3-bisphosphoglycerate-dependent phosphoglycerate mutase |
| group_9547 | gpx1 | Hydroperoxy fatty acid reductase gpx1 |
| group_9510 | gspS2 | Pilotin AspS 2 |
| group_9955 | gstB | Glutathione S-transferase GST-6.0 |
| group_10162 | hcr | NADH oxidoreductase HCR |
| hlyA_2 | hlyA_2 | Hemolysin |
| hlyU | hlyU | Transcriptional activator HlyU |
| group_9765 | hmp_1 | Flavohemoprotein |
| group_9627 | hmuU_1 | Hemin transport system permease protein HmuU |
| group_9626 | hmuV_1 | Hemin import ATP-binding protein HmuV |
| group_10093 | hpd | 4-hydroxyphenylpyruvate dioxygenase |
| group_10106 | inhA_2 | Isonitrile hydratase |
| intA_3 | intA_3 | Prophage integrase IntA |
| iolG | iolG | Inositol 2-dehydrogenase/D-chiro-inositol 3-dehydrogenase |
| group_269 | ISSde3 | IS30 family transposase ISSde3 |
| group_270 | ISSde3 | IS30 family transposase ISSde3 |
| kdgK_3 | kdgK_3 | 2-dehydro-3-deoxygluconokinase |
| group_10182 | kdpD | Sensor protein KdpD |
| kdsA_2 | kdsA_2 | 2-dehydro-3-deoxyphosphooctonate aldolase |
| kdsB_2 | kdsB_2 | 3-deoxy-manno-octulosonate cytidylyltransferase |
| kpsF | kpsF | Arabinose 5-phosphate isomerase KpsF |
| group_9900 | lepB_2 | Signal peptidase I |
| group_10025 | lhgD | L-2-hydroxyglutarate dehydrogenase |
| group_9814 | lig | DNA ligase |
| livF | livF | High-affinity branched-chain amino acid transport ATP-binding protein LivF |
| lptB_2 | lptB_2 | Lipopolysaccharide export system ATP-binding protein LptB |
| group_9528 | lutC | Lactate utilization protein C |
| maa | maa | Maltose O-acetyltransferase |
| group_10109 | macA | Macrolide export protein MacA |
| group_10111 | macB_1 | Macrolide export ATP-binding/permease protein MacB |
| group_10112 | macB_2 | Macrolide export ATP-binding/permease protein MacB |
| group_9818 | mak | Fructokinase |
| malX_2 | malX_2 | Maltose/maltodextrin-binding protein |
| mddA | mddA | Methanethiol S-methyltransferase |
| group_9517 | mdtA_1 | Multidrug resistance protein MdtA |
| mdtL_2 | mdtL_2 | Multidrug resistance protein MdtL |
| group_9991 | mdtN | Multidrug resistance protein MdtN |
| menC_1 | menC_1 | o-succinylbenzoate synthase |
| group_9739 | menH_2 | 2-succinyl-6-hydroxy-2,4-cyclohexadiene-1-carboxylate synthase |
| group_10031 | mepM_1 | Murein DD-endopeptidase MepM |
| group_10239 | metL | Bifunctional aspartokinase/homoserine dehydrogenase 2 |
| group_9653 | mltF_2 | Membrane-bound lytic murein transglycosylase F |
| mnaA | mnaA | UDP-N-acetylglucosamine 2-epimerase |
| group_9742 | mnmC | tRNA 5-methylaminomethyl-2-thiouridine biosynthesis bifunctional protein MnmC |
| group_9890 | mntH | Divalent metal cation transporter MntH |
| group_9837 | modE | DNA-binding transcriptional dual regulator ModE |
| group_10146 | mprA_2 | Response regulator MprA |
| group_10053 | mrdA_2 | Peptidoglycan D,D-transpeptidase MrdA |
| group_10107 | mreB_1 | Cell shape-determining protein MreB |
| group_2175 | mscM_1 | Miniconductance mechanosensitive channel MscM |
| mshA_1 | mshA_1 | D-inositol-3-phosphate glycosyltransferase |
| group_9722 | mshA_1 | D-inositol-3-phosphate glycosyltransferase |
| group_9776 | mshA_2 | D-inositol-3-phosphate glycosyltransferase |
| group_10132 | msrA3 | Peptide methionine sulfoxide reductase MsrA 3 |
| group_9733 | msrB | Peptide methionine sulfoxide reductase MsrB |
| group_9859 | msrQ | Protein-methionine-sulfoxide reductase heme-binding subunit MsrQ |
| group_9796 | murB | UDP-N-acetylenolpyruvoylglucosamine reductase |
| group_9554 | murE | UDP-N-acetylmuramoyl-L-alanyl-D-glutamate--2,6-diaminopimelate ligase |
| group_9766 | mutL | DNA mismatch repair protein MutL |
| group_9882 | nadD | nicotinate-nucleotide adenylyltransferase |
| group_4042 | nagA | N-acetylglucosamine-6-phosphate deacetylase |
| group_9842 | nagL | Maleylpyruvate isomerase |
| group_9684 | nudC_2 | NADH pyrophosphatase |
| nudF_1 | nudF_1 | ADP-ribose pyrophosphatase |
| group_10175 | ogt_2 | Methylated-DNA--protein-cysteine methyltransferase |
| group_10029 | ompA_6 | Outer membrane protein A |
| group_9498 | ompH | Porin-like protein H |
| group_10036 | opgE | Phosphoethanolamine transferase OpgE |
| group_9594 | outN | Type II secretion system protein N |
| group_9929 | paaF | 2,3-dehydroadipyl-CoA hydratase |
| group_9912 | pac | Penicillin acylase |
| group_9976 | pal_2 | Peptidoglycan-associated lipoprotein |
| group_9774 | pal_5 | Peptidoglycan-associated lipoprotein |
| group_9625 | parB | putative chromosome-partitioning protein ParB |
| group_9697 | pdeG | putative cyclic di-GMP phosphodiesterase PdeG |
| group_9810 | pepT_2 | Peptidase T |
| group_9835 | pgrR_2 | HTH-type transcriptional regulator PgrR |
| group_9971 | phrB | (6-4) photolyase |
| phrB_2 | phrB_2 | Deoxyribodipyrimidine photo-lyase |
| group_9553 | pilA | Fimbrial protein |
| group_10142 | pip | Proline iminopeptidase |
| group_9999 | PLA1A | Putative phospholipase A1 |
| group_7698 | pldA | Phospholipase A1 |
| group_9970 | pldB_1 | Lysophospholipase L2 |
| group_10253 | pldB_2 | Lysophospholipase L2 |
| group_10097 | plsC_2 | 1-acyl-sn-glycerol-3-phosphate acyltransferase |
| group_10083 | potD_2 | Spermidine/putrescine-binding periplasmic protein |
| group_10209 | pse4 | Beta-lactamase PSE-4 |
| group_9808 | puuR | HTH-type transcriptional regulator PuuR |
| group_9674 | pxpA3 | 5-oxoprolinase subunit A 3 |
| group_9673 | pxpB | 5-oxoprolinase subunit B |
| group_9672 | pxpC | 5-oxoprolinase subunit C |
| group_9819 | rbgA | Ribosome biogenesis GTPase A |
| group_9569 | rcsC_2 | Sensor histidine kinase RcsC |
| group_9772 | rcsC_5 | Sensor histidine kinase RcsC |
| group_10052 | rcsC_6 | Sensor histidine kinase RcsC |
| rcsC_8 | rcsC_8 | Sensor histidine kinase RcsC |
| group_10095 | resA | Thiol-disulfide oxidoreductase ResA |
| group_9753 | rfbC | dTDP-4-dehydrorhamnose 3,5-epimerase |
| group_9755 | rffG_2 | dTDP-glucose 4,6-dehydratase 2 |
| group_9754 | rffH | Glucose-1-phosphate thymidylyltransferase 2 |
| group_10065 | rhaS_2 | HTH-type transcriptional activator RhaS |
| group_10070 | rlpA_2 | Endolytic peptidoglycan transglycosylase RlpA |
| rpoD_2 | rpoD_2 | RNA polymerase sigma factor RpoD |
| group_9889 | rpoS_2 | RNA polymerase sigma factor RpoS |
| group_10017 | rpoS_3 | RNA polymerase sigma factor RpoS |
| group_10273 | RRAAH_MYCLE | Putative 4-hydroxy-4-methyl-2-oxoglutarate aldolase |
| group_10163 | sadA | Autotransporter adhesin SadA |
| group_9515 | sapB | Putrescine export system permease protein SapB |
| group_9923 | sasA_6 | Adaptive-response sensory-kinase SasA |
| group_9720 | sdaC_4 | Serine transporter |
| group_9843 | Sez_1297 | Phosphorylated carbohydrates phosphatase |
| sfaA | sfaA | S-fimbrial protein subunit SfaA |
| group_560 | sigM | ECF RNA polymerase sigma factor SigM |
| sigW | sigW | ECF RNA polymerase sigma factor SigW |
| spnR | spnR | dTDP-4-dehydro-2,3,6-trideoxy-D-glucose 4-aminotransferase |
| sseA | sseA | 3-mercaptopyruvate sulfurtransferase |
| group_9784 | syrM1 | HTH-type transcriptional regulator SyrM 1 |
| group_10118 | thiD | Hydroxymethylpyrimidine/phosphomethylpyrimidine kinase |
| group_9590 | thiF | Sulfur carrier protein ThiS adenylyltransferase |
| group_9614 | tilS | tRNA(Ile)-lysidine synthase |
| group_9785 | tmcA | tRNA(Met) cytidine acetyltransferase TmcA |
| group_10110 | tolC_2 | Outer membrane protein TolC |
| group_10150 | tolR_3 | Tol-Pal system protein TolR |
| group_10188 | tpx_1 | Thiol peroxidase |
| group_10189 | tpx_2 | Thiol peroxidase |
| group_9782 | tuaC | Putative teichuronic acid biosynthesis glycosyltransferase TuaC |
| group_9659 | tyrP_2 | Tyrosine-specific transport protein |
| ureA | ureA | Urease subunit gamma |
| ureB | ureB | Urease subunit beta |
| ureC | ureC | Urease subunit alpha |
| ureD | ureD | Urease accessory protein UreD |
| ureE | ureE | Urease accessory protein UreE |
| ureF | ureF | Urease accessory protein UreF |
| ureG_1 | ureG_1 | Urease accessory protein UreG |
| group_10205 | uvrB_2 | UvrABC system protein B |
| group_9703 | volA_1 | Lysophospholipase VolA |
| wbnH | wbnH | O-antigen biosynthesis glycosyltransferase WbnH |
| group_9746 | wecA | Undecaprenyl-phosphate alpha-N-acetylglucosaminyl 1-phosphate transferase |
| wecB | wecB | UDP-N-acetylglucosamine 2-epimerase |
| group_9892 | xerC_2 | Tyrosine recombinase XerC |
| group_9914 | xerC_2 | Tyrosine recombinase XerC |
| group_10158 | yabJ_1 | 2-iminobutanoate/2-iminopropanoate deaminase |
| group_9853 | ybbJ | Inner membrane protein YbbJ |
| group_9663 | ybdG | Miniconductance mechanosensitive channel YbdG |
| group_9500 | ybfF | Esterase YbfF |
| group_9867 | ycaC | putative hydrolase YcaC |
| group_9638 | ycbB_2 | putative L,D-transpeptidase YcbB |
| group_9888 | yceB | putative lipoprotein YceB |
| yceJ_1 | yceJ_1 | Cytochrome b561 |
| group_10221 | yceJ_2 | Cytochrome b561 |
| group_9897 | yceM | Putative oxidoreductase YceM |
| ycfH_2 | ycfH_2 | putative metal-dependent hydrolase YcfH |
| yddE | yddE | putative isomerase YddE |
| group_9734 | yeaD | Putative glucose-6-phosphate 1-epimerase |
| group_10032 | yedI | Inner membrane protein YedI |
| group_9543 | yeeA_1 | Inner membrane protein YeeA |
| group_10028 | yeeN | putative transcriptional regulatory protein YeeN |
| group_9516 | yeeZ | Protein YeeZ |
| group_9529 | yhjX_2 | putative MFS-type transporter YhjX |
| group_9524 | yjcS | Putative alkyl/aryl-sulfatase YjcS |
| group_10105 | yjgH | RutC family protein YjgH |
| group_9963 | yjjG | Pyrimidine 5'-nucleotidase YjjG |
| group_10270 | yjjV | putative metal-dependent hydrolase YjjV |
| group_9639 | yjjW | Putative glycyl-radical enzyme activating enzyme YjjW |
| group_10113 | yknY | putative ABC transporter ATP-binding protein YknY |
| group_9685 | ynjE | Thiosulfate sulfurtransferase YnjE |
| group_9994 | ypdF | Aminopeptidase YpdF |
| group_10156 | yqaB_2 | Fructose-1-phosphate phosphatase YqaB |
| yqjZ_2 | yqjZ_2 | putative protein YqjZ |
| group_10144 | ysnE | putative N-acetyltransferase YsnE |
| yvbK | yvbK | putative N-acetyltransferase YvbK |
| group_10119 | ywrO_1 | General stress protein 14 |
| group_9913 | yxeI_2 | putative protein YxeI |
| group_9568 | zapC | Cell division protein ZapC |
| group_9630 | zntR | HTH-type transcriptional regulator ZntR |
| group_10152 |  | Pentapeptide repeat protein |

**Table S3.** Genes present in plasmid sequences of the *Photobacterium* strains indicated and their copy number detected. Reference sequences from NCBI included in the analysis are indicated with a *.

| **gene** | **ajapo4.1*** | **ajapo5.5** | **ajapo8.2** | **ATCC25521** | **Ik8.1** | **Ik8.2** | **Kume1.2** | **Kume1.3** | **JS01*** | **lnuch19.1** | **lsplen1.1** |
| --- | --- | --- | --- | --- | --- | --- | --- | --- | --- | --- | --- |
| accD | 0 | 0 | 0 | 0 | 0 | 0 | 0 | 0 | 0 | 0 | 0 |
| Acetyltransferase | 0 | 0 | 0 | 0 | 0 | 0 | 0 | 0 | 0 | 0 | 1 |
| adhB | 0 | 0 | 0 | 0 | 1 | 1 | 0 | 0 | 1 | 2 | 1 |
| aguA | 0 | 0 | 0 | 0 | 0 | 0 | 0 | 0 | 0 | 0 | 0 |
| argO | 0 | 0 | 0 | 0 | 0 | 0 | 0 | 0 | 1 | 0 | 1 |
| bin3 | 1 | 1 | 1 | 0 | 0 | 1 | 0 | 0 | 1 | 1 | 1 |
| bioF | 0 | 0 | 0 | 0 | 0 | 0 | 0 | 0 | 0 | 0 | 0 |
| btr | 0 | 0 | 0 | 0 | 0 | 0 | 0 | 0 | 0 | 0 | 0 |
| btuD | 0 | 0 | 0 | 0 | 0 | 0 | 0 | 0 | 0 | 0 | 0 |
| cas1 | 0 | 0 | 0 | 0 | 0 | 0 | 0 | 0 | 0 | 0 | 0 |
| cas2 | 0 | 0 | 0 | 0 | 0 | 0 | 0 | 0 | 0 | 0 | 0 |
| cobS | 0 | 0 | 0 | 0 | 0 | 0 | 0 | 0 | 0 | 0 | 0 |
| copR | 0 | 0 | 0 | 0 | 0 | 0 | 0 | 0 | 0 | 0 | 0 |
| cpdA | 0 | 0 | 0 | 0 | 0 | 0 | 0 | 0 | 0 | 0 | 0 |
| cspG | 0 | 0 | 0 | 0 | 0 | 0 | 0 | 0 | 0 | 1 | 0 |
| cyaB | 0 | 0 | 0 | 0 | 0 | 0 | 0 | 0 | 0 | 0 | 0 |
| cyt1Aa | 0 | 0 | 0 | 0 | 0 | 0 | 0 | 0 | 1 | 0 | 0 |
| dapH | 0 | 0 | 0 | 0 | 0 | 0 | 0 | 0 | 0 | 0 | 1 |
| ddl | 0 | 0 | 0 | 0 | 0 | 0 | 0 | 0 | 0 | 0 | 1 |
| degP | 0 | 0 | 0 | 0 | 0 | 0 | 0 | 0 | 0 | 0 | 0 |
| degQ | 0 | 0 | 0 | 0 | 0 | 0 | 0 | 0 | 0 | 0 | 0 |
| dinB | 0 | 0 | 0 | 2 | 0 | 0 | 0 | 0 | 0 | 0 | 0 |
| dnaG | 0 | 0 | 0 | 0 | 0 | 0 | 0 | 0 | 0 | 0 | 1 |
| dns | 1 | 1 | 0 | 0 | 0 | 0 | 2 | 0 | 0 | 1 | 2 |
| dosP | 0 | 0 | 0 | 0 | 0 | 0 | 0 | 0 | 0 | 0 | 0 |
| echA8 | 0 | 0 | 0 | 0 | 0 | 0 | 0 | 0 | 0 | 0 | 0 |
| elaA | 0 | 0 | 0 | 0 | 0 | 0 | 0 | 0 | 0 | 0 | 1 |
| elmMIII | 0 | 0 | 0 | 0 | 0 | 0 | 0 | 0 | 0 | 0 | 1 |
| emrE | 0 | 0 | 0 | 0 | 0 | 0 | 0 | 0 | 0 | 1 | 1 |
| epsE | 0 | 0 | 0 | 0 | 0 | 0 | 0 | 0 | 0 | 0 | 1 |
| exsA_1 | 0 | 0 | 0 | 0 | 0 | 0 | 0 | 0 | 0 | 0 | 0 |
| Extracellular serine proteinase | 0 | 0 | 0 | 0 | 0 | 0 | 0 | 0 | 0 | 0 | 0 |
| Ferredoxin--NADP reductase | 0 | 0 | 0 | 0 | 0 | 0 | 0 | 0 | 0 | 0 | 0 |
| ghrA | 0 | 0 | 0 | 0 | 0 | 0 | 0 | 0 | 0 | 0 | 1 |
| gmk | 0 | 0 | 0 | 0 | 0 | 0 | 0 | 0 | 0 | 0 | 0 |
| groL | 0 | 0 | 0 | 0 | 0 | 0 | 0 | 0 | 0 | 0 | 1 |
| groS | 0 | 0 | 0 | 0 | 0 | 0 | 0 | 0 | 0 | 0 | 1 |
| gst | 0 | 0 | 0 | 0 | 0 | 0 | 0 | 0 | 0 | 0 | 1 |
| hapE | 0 | 0 | 0 | 0 | 0 | 0 | 0 | 0 | 0 | 0 | 0 |
| higA1 | 0 | 0 | 0 | 0 | 0 | 0 | 0 | 0 | 0 | 0 | 0 |
| hipA | 0 | 0 | 0 | 0 | 0 | 1 | 0 | 0 | 0 | 0 | 0 |
| hisC2 | 0 | 0 | 0 | 0 | 0 | 0 | 0 | 0 | 0 | 0 | 1 |
| hns | 0 | 0 | 0 | 0 | 0 | 0 | 0 | 0 | 0 | 1 | 1 |
| hsdR | 0 | 0 | 0 | 0 | 0 | 0 | 0 | 0 | 0 | 0 | 0 |
| htrC | 0 | 0 | 0 | 0 | 0 | 0 | 0 | 0 | 0 | 0 | 0 |
| hup | 0 | 0 | 0 | 0 | 0 | 0 | 0 | 0 | 0 | 0 | 1 |
| hxpA | 0 | 0 | 0 | 0 | 0 | 0 | 0 | 0 | 0 | 0 | 0 |
| ilvE | 0 | 0 | 0 | 0 | 0 | 0 | 0 | 0 | 0 | 0 | 1 |
| imm | 0 | 0 | 0 | 0 | 0 | 0 | 0 | 0 | 0 | 0 | 0 |
| invA | 0 | 0 | 0 | 0 | 0 | 0 | 0 | 0 | 0 | 0 | 0 |
| IS1 family transposase ISPda1 | 0 | 0 | 0 | 0 | 0 | 0 | 0 | 0 | 0 | 0 | 0 |
| IS110 family transposase ISCps1 | 0 | 0 | 0 | 0 | 0 | 0 | 0 | 0 | 0 | 0 | 1 |
| IS110 family transposase ISSde4 | 1 | 0 | 1 | 0 | 0 | 0 | 0 | 0 | 0 | 0 | 0 |
| IS110 family transposase ISSpi6 | 0 | 0 | 0 | 0 | 0 | 0 | 0 | 0 | 0 | 0 | 0 |
| IS200/IS605 family transposase ISEc42 | 0 | 0 | 0 | 0 | 0 | 0 | 0 | 0 | 0 | 0 | 0 |
| IS200/IS605 family transposase ISEc46 | 0 | 0 | 0 | 0 | 0 | 0 | 0 | 0 | 0 | 0 | 0 |
| IS21 family transposase ISVch3 | 0 | 0 | 0 | 0 | 0 | 0 | 0 | 0 | 0 | 0 | 0 |
| IS3 family transposase ISBps2 | 0 | 0 | 0 | 0 | 0 | 0 | 0 | 0 | 0 | 0 | 0 |
| IS3 family transposase ISThsp4 | 0 | 0 | 0 | 0 | 0 | 0 | 0 | 0 | 0 | 0 | 0 |
| IS3 family transposase ISVch4 | 0 | 0 | 0 | 0 | 0 | 0 | 0 | 0 | 0 | 0 | 0 |
| IS30 family transposase ISSde3 | 0 | 0 | 0 | 0 | 0 | 0 | 0 | 0 | 0 | 0 | 0 |
| IS4 family transposase ISAzvi5 | 0 | 0 | 0 | 0 | 0 | 0 | 0 | 0 | 0 | 0 | 0 |
| IS4 family transposase ISCro6 | 0 | 0 | 0 | 0 | 0 | 0 | 0 | 0 | 0 | 0 | 0 |
| IS4 family transposase ISPcc6 | 0 | 0 | 0 | 0 | 0 | 0 | 0 | 0 | 0 | 0 | 0 |
| IS4 family transposase ISVa18 | 0 | 0 | 0 | 0 | 0 | 0 | 0 | 0 | 0 | 0 | 0 |
| IS4 family transposase ISVsa5 | 0 | 0 | 0 | 0 | 0 | 0 | 0 | 0 | 0 | 0 | 0 |
| IS5 family transposase ISVha3 | 0 | 0 | 0 | 0 | 0 | 0 | 0 | 0 | 0 | 0 | 0 |
| IS6 family transposase ISPpr9 | 0 | 0 | 1 | 0 | 0 | 0 | 0 | 0 | 2 | 6 | 11 |
| IS630 family transposase ISVa15 | 0 | 0 | 1 | 0 | 0 | 0 | 0 | 0 | 0 | 0 | 1 |
| IS66 family transposase ISVa11 | 0 | 0 | 0 | 0 | 0 | 0 | 0 | 0 | 0 | 1 | 0 |
| IS66 family transposase ISVa5 | 0 | 0 | 0 | 0 | 0 | 0 | 0 | 0 | 0 | 0 | 0 |
| IS66 family transposase ISVa9 | 0 | 0 | 0 | 0 | 0 | 0 | 0 | 0 | 0 | 0 | 0 |
| IS91 family transposase ISShvi3 | 0 | 0 | 0 | 0 | 0 | 0 | 0 | 0 | 0 | 0 | 1 |
| IS91 family transposase ISVsa9 | 0 | 0 | 0 | 0 | 0 | 0 | 0 | 0 | 0 | 0 | 1 |
| ISAs1 family transposase ISVha1 | 0 | 0 | 0 | 0 | 0 | 0 | 0 | 0 | 0 | 0 | 0 |
| Isatin hydrolase | 0 | 0 | 0 | 0 | 0 | 0 | 0 | 0 | 0 | 0 | 1 |
| ISNCY family transposase ISLad2 | 0 | 0 | 0 | 0 | 0 | 0 | 0 | 0 | 0 | 0 | 0 |
| ISNCY family transposase ISLad2 | 0 | 0 | 0 | 0 | 0 | 0 | 0 | 0 | 0 | 0 | 0 |
| ISNCY family transposase ISVisp7 | 0 | 0 | 0 | 0 | 0 | 0 | 0 | 0 | 0 | 0 | 0 |
| kynA | 0 | 0 | 0 | 0 | 0 | 0 | 0 | 0 | 0 | 0 | 1 |
| kynu | 0 | 0 | 0 | 0 | 0 | 0 | 0 | 0 | 0 | 0 | 1 |
| lagD | 0 | 0 | 0 | 0 | 1 | 1 | 0 | 0 | 0 | 1 | 0 |
| lcdH | 0 | 0 | 0 | 0 | 0 | 0 | 0 | 0 | 0 | 0 | 2 |
| lgrD | 0 | 0 | 0 | 0 | 0 | 0 | 0 | 0 | 0 | 0 | 0 |
| lysO | 0 | 0 | 0 | 0 | 0 | 0 | 0 | 0 | 0 | 0 | 0 |
| mhpA | 0 | 0 | 0 | 0 | 0 | 0 | 0 | 0 | 0 | 0 | 1 |
| moaA | 0 | 0 | 0 | 0 | 0 | 0 | 0 | 0 | 0 | 0 | 0 |
| mtcA2 | 0 | 0 | 0 | 0 | 0 | 0 | 0 | 0 | 0 | 0 | 0 |
| N-carbamoyl-D-amino acid hydrolase | 0 | 0 | 0 | 0 | 0 | 0 | 0 | 0 | 0 | 0 | 0 |
| nimR | 2 | 0 | 2 | 0 | 0 | 0 | 0 | 0 | 0 | 0 | 0 |
| nrgA | 0 | 0 | 0 | 0 | 0 | 0 | 0 | 0 | 0 | 0 | 0 |
| ntdA | 0 | 0 | 0 | 0 | 0 | 0 | 0 | 0 | 0 | 0 | 0 |
| ntdB | 0 | 0 | 0 | 0 | 0 | 0 | 0 | 0 | 0 | 0 | 0 |
| ntdC | 0 | 0 | 0 | 0 | 0 | 0 | 0 | 0 | 0 | 0 | 0 |
| nucM | 0 | 0 | 0 | 0 | 0 | 0 | 0 | 0 | 0 | 0 | 0 |
| ompA | 0 | 0 | 0 | 0 | 0 | 0 | 0 | 0 | 0 | 0 | 1 |
| parA | 2 | 0 | 0 | 0 | 0 | 0 | 0 | 0 | 0 | 0 | 0 |
| pdeG | 0 | 0 | 0 | 0 | 0 | 0 | 0 | 0 | 0 | 0 | 0 |
| pgrR | 0 | 0 | 0 | 0 | 0 | 0 | 0 | 0 | 0 | 0 | 1 |
| pinR | 0 | 0 | 0 | 0 | 1 | 0 | 2 | 1 | 0 | 0 | 3 |
| pndA | 0 | 0 | 0 | 0 | 0 | 0 | 0 | 0 | 0 | 0 | 1 |
| pndA | 0 | 0 | 0 | 0 | 0 | 0 | 0 | 0 | 0 | 0 | 0 |
| proQ | 0 | 0 | 0 | 0 | 0 | 0 | 0 | 0 | 0 | 0 | 0 |
| prtR | 0 | 0 | 0 | 0 | 0 | 0 | 0 | 0 | 0 | 0 | 1 |
| pscF | 0 | 0 | 0 | 0 | 0 | 0 | 0 | 0 | 0 | 0 | 0 |
| ptlH | 0 | 0 | 0 | 0 | 0 | 0 | 0 | 0 | 0 | 0 | 1 |
| Putative nickel-responsive regulator | 0 | 0 | 0 | 0 | 0 | 0 | 0 | 0 | 0 | 0 | 0 |
| putative oxidoreductase | 0 | 0 | 0 | 0 | 0 | 0 | 0 | 0 | 0 | 0 | 1 |
| queuosine precursor transporter | 0 | 0 | 0 | 0 | 0 | 0 | 0 | 0 | 0 | 1 | 0 |
| ramA | 0 | 0 | 0 | 0 | 0 | 0 | 0 | 0 | 1 | 0 | 0 |
| ramC | 0 | 0 | 0 | 0 | 0 | 0 | 0 | 0 | 0 | 1 | 0 |
| rclC | 0 | 0 | 0 | 0 | 0 | 0 | 0 | 0 | 0 | 0 | 1 |
| recX | 0 | 0 | 0 | 0 | 0 | 0 | 0 | 0 | 0 | 0 | 0 |
| rep | 0 | 0 | 0 | 0 | 0 | 0 | 0 | 0 | 0 | 0 | 0 |
| repA | 1 | 1 | 1 | 0 | 0 | 0 | 2 | 1 | 0 | 1 | 1 |
| rhaR | 0 | 0 | 0 | 0 | 0 | 0 | 0 | 0 | 0 | 1 | 0 |
| rhaS | 0 | 0 | 0 | 0 | 0 | 0 | 0 | 0 | 0 | 0 | 1 |
| rhsC | 0 | 0 | 0 | 0 | 0 | 0 | 0 | 0 | 0 | 1 | 0 |
| ribD | 0 | 0 | 0 | 0 | 0 | 0 | 0 | 0 | 0 | 0 | 0 |
| ribF | 0 | 0 | 0 | 0 | 0 | 0 | 0 | 0 | 0 | 1 | 0 |
| rizA | 0 | 0 | 0 | 0 | 0 | 0 | 0 | 0 | 0 | 0 | 1 |
| rutB | 0 | 0 | 0 | 0 | 0 | 0 | 0 | 0 | 1 | 0 | 1 |
| sasA | 0 | 0 | 0 | 0 | 0 | 0 | 0 | 0 | 0 | 0 | 0 |
| sctC | 0 | 0 | 0 | 0 | 0 | 0 | 0 | 0 | 0 | 0 | 0 |
| slyA | 0 | 0 | 0 | 0 | 1 | 1 | 0 | 0 | 0 | 0 | 0 |
| smc | 0 | 0 | 0 | 0 | 0 | 1 | 0 | 0 | 0 | 0 | 0 |
| sodC | 0 | 0 | 0 | 0 | 0 | 0 | 0 | 0 | 0 | 0 | 1 |
| soj | 0 | 0 | 0 | 0 | 0 | 0 | 0 | 0 | 0 | 0 | 0 |
| soxR | 0 | 0 | 0 | 0 | 0 | 0 | 0 | 0 | 0 | 0 | 0 |
| spaR | 0 | 0 | 0 | 0 | 0 | 0 | 0 | 0 | 0 | 0 | 0 |
| tcpE | 0 | 0 | 0 | 0 | 0 | 0 | 0 | 0 | 0 | 0 | 0 |
| TDO2 | 0 | 0 | 0 | 0 | 0 | 0 | 0 | 0 | 0 | 0 | 1 |
| tetA | 0 | 0 | 0 | 0 | 0 | 0 | 0 | 0 | 0 | 0 | 1 |
| tfdR | 0 | 0 | 0 | 0 | 0 | 0 | 0 | 0 | 1 | 0 | 0 |
| tmcA | 2 | 0 | 0 | 0 | 0 | 0 | 0 | 0 | 0 | 0 | 0 |
| Tn3 family transposase ISSod9 | 0 | 0 | 0 | 0 | 0 | 0 | 0 | 0 | 0 | 0 | 0 |
| Tn3 family transposase ISXc4 | 0 | 0 | 0 | 0 | 0 | 0 | 0 | 0 | 0 | 0 | 0 |
| tnpR | 1 | 1 | 1 | 1 | 1 | 0 | 2 | 1 | 1 | 1 | 2 |
| topA | 0 | 0 | 0 | 0 | 0 | 0 | 0 | 0 | 0 | 0 | 0 |
| topB | 0 | 0 | 0 | 0 | 0 | 0 | 0 | 0 | 0 | 0 | 1 |
| traC | 0 | 0 | 0 | 0 | 0 | 0 | 0 | 0 | 0 | 1 | 2 |
| traD | 0 | 0 | 0 | 0 | 0 | 0 | 0 | 0 | 0 | 0 | 1 |
| traN | 0 | 0 | 0 | 0 | 0 | 0 | 0 | 0 | 0 | 0 | 1 |
| tyeA | 0 | 0 | 0 | 0 | 0 | 0 | 0 | 0 | 0 | 0 | 0 |
| virB | 0 | 0 | 0 | 0 | 0 | 0 | 0 | 0 | 0 | 1 | 0 |
| virB1 | 0 | 0 | 0 | 0 | 0 | 0 | 0 | 0 | 0 | 0 | 0 |
| virB4 | 0 | 0 | 0 | 0 | 0 | 0 | 0 | 0 | 0 | 0 | 0 |
| xerC | 0 | 0 | 0 | 0 | 0 | 0 | 0 | 0 | 0 | 1 | 0 |
| xerD | 0 | 0 | 0 | 0 | 0 | 0 | 0 | 0 | 0 | 0 | 1 |
| yedJ | 0 | 0 | 0 | 0 | 0 | 0 | 0 | 0 | 0 | 0 | 0 |
| yegI | 0 | 0 | 0 | 0 | 0 | 0 | 0 | 0 | 0 | 0 | 0 |
| ygiW | 0 | 0 | 0 | 0 | 0 | 0 | 0 | 0 | 0 | 0 | 0 |
| yhcG | 0 | 0 | 0 | 0 | 0 | 0 | 0 | 0 | 0 | 1 | 0 |
| yiaC | 0 | 0 | 0 | 0 | 0 | 0 | 0 | 0 | 0 | 0 | 0 |
| yigB | 0 | 0 | 0 | 0 | 0 | 0 | 0 | 0 | 0 | 1 | 0 |
| yopB | 0 | 0 | 0 | 0 | 0 | 0 | 0 | 0 | 0 | 0 | 0 |
| yopN | 0 | 0 | 0 | 0 | 0 | 0 | 0 | 0 | 0 | 0 | 0 |
| yraJ | 0 | 0 | 0 | 0 | 0 | 0 | 0 | 0 | 0 | 0 | 0 |
| yscD | 0 | 0 | 0 | 0 | 0 | 0 | 0 | 0 | 0 | 0 | 0 |
| yscG | 0 | 0 | 0 | 0 | 0 | 0 | 0 | 0 | 0 | 0 | 0 |
| yscJ | 0 | 0 | 0 | 0 | 0 | 0 | 0 | 0 | 0 | 0 | 0 |
| yscN | 0 | 0 | 0 | 0 | 0 | 0 | 0 | 0 | 0 | 0 | 0 |
| yscU | 0 | 0 | 0 | 0 | 0 | 0 | 0 | 0 | 0 | 0 | 0 |
| **gene** | **StJ4.21** | **ajapo8.1** | **ljone10.1** | **LN-1a** | **lequu1.1** | **LF-1a** | **LN-I.1** | **LR-VIII.1** | **svers1.1*** | **ahane1.5** | **calba1.1** |
| accD | 0 | 0 | 1 | 0 | 0 | 0 | 0 | 0 | 0 | 0 | 0 |
| Acetyltransferase | 0 | 0 | 0 | 0 | 0 | 0 | 0 | 0 | 0 | 0 | 0 |
| adhB | 0 | 0 | 0 | 0 | 1 | 1 | 0 | 0 | 0 | 1 | 0 |
| aguA | 0 | 0 | 0 | 0 | 0 | 0 | 0 | 0 | 0 | 1 | 0 |
| argO | 0 | 0 | 0 | 0 | 1 | 1 | 0 | 0 | 0 | 0 | 0 |
| bin3 | 0 | 2 | 0 | 0 | 1 | 0 | 4 | 0 | 1 | 0 | 0 |
| bioF | 0 | 0 | 0 | 0 | 1 | 0 | 0 | 0 | 0 | 0 | 0 |
| btr | 0 | 0 | 1 | 0 | 0 | 0 | 0 | 0 | 0 | 0 | 0 |
| btuD | 0 | 0 | 1 | 0 | 0 | 0 | 0 | 0 | 0 | 0 | 0 |
| cas1 | 0 | 0 | 0 | 0 | 0 | 0 | 0 | 0 | 0 | 0 | 1 |
| cas2 | 0 | 0 | 0 | 0 | 0 | 0 | 0 | 0 | 0 | 0 | 1 |
| cobS | 1 | 0 | 0 | 0 | 0 | 0 | 0 | 0 | 0 | 0 | 0 |
| copR | 0 | 0 | 0 | 0 | 0 | 0 | 1 | 0 | 0 | 0 | 0 |
| cpdA | 0 | 0 | 0 | 1 | 0 | 0 | 0 | 0 | 0 | 1 | 0 |
| cspG | 0 | 0 | 0 | 1 | 0 | 0 | 0 | 0 | 0 | 2 | 0 |
| cyaB | 0 | 0 | 0 | 0 | 0 | 0 | 0 | 0 | 0 | 0 | 1 |
| cyt1Aa | 0 | 0 | 0 | 0 | 0 | 0 | 0 | 0 | 0 | 0 | 0 |
| dapH | 0 | 0 | 0 | 0 | 0 | 0 | 1 | 1 | 0 | 0 | 0 |
| ddl | 0 | 0 | 0 | 0 | 0 | 0 | 0 | 0 | 0 | 0 | 0 |
| degP | 0 | 0 | 2 | 0 | 0 | 0 | 1 | 0 | 0 | 0 | 0 |
| degQ | 0 | 0 | 1 | 0 | 0 | 0 | 0 | 0 | 0 | 0 | 0 |
| dinB | 0 | 0 | 0 | 0 | 0 | 0 | 0 | 0 | 0 | 0 | 0 |
| dnaG | 0 | 0 | 0 | 0 | 0 | 0 | 0 | 0 | 0 | 0 | 0 |
| dns | 0 | 0 | 0 | 0 | 0 | 1 | 0 | 1 | 1 | 1 | 0 |
| dosP | 0 | 0 | 0 | 0 | 0 | 0 | 0 | 0 | 0 | 1 | 0 |
| echA8 | 0 | 0 | 0 | 0 | 0 | 0 | 1 | 0 | 0 | 0 | 0 |
| elaA | 0 | 0 | 0 | 0 | 0 | 0 | 0 | 0 | 0 | 0 | 0 |
| elmMIII | 0 | 0 | 0 | 0 | 0 | 0 | 0 | 0 | 0 | 0 | 0 |
| emrE | 0 | 0 | 0 | 0 | 0 | 0 | 0 | 0 | 0 | 0 | 0 |
| epsE | 0 | 0 | 0 | 0 | 0 | 0 | 0 | 2 | 0 | 0 | 0 |
| exsA_1 | 2 | 0 | 0 | 0 | 0 | 0 | 0 | 0 | 0 | 0 | 0 |
| Extracellular serine proteinase | 0 | 0 | 1 | 0 | 0 | 0 | 0 | 0 | 0 | 0 | 0 |
| Ferredoxin--NADP reductase | 0 | 0 | 0 | 0 | 0 | 0 | 1 | 0 | 0 | 0 | 0 |
| ghrA | 0 | 0 | 0 | 0 | 0 | 0 | 0 | 0 | 0 | 0 | 0 |
| gmk | 0 | 0 | 0 | 0 | 0 | 0 | 1 | 0 | 0 | 0 | 0 |
| groL | 0 | 0 | 0 | 0 | 0 | 0 | 1 | 0 | 0 | 0 | 0 |
| groS | 0 | 0 | 0 | 0 | 0 | 0 | 1 | 0 | 0 | 0 | 0 |
| gst | 0 | 0 | 0 | 0 | 0 | 0 | 0 | 0 | 0 | 0 | 0 |
| hapE | 0 | 0 | 0 | 0 | 0 | 0 | 0 | 1 | 0 | 0 | 0 |
| higA1 | 0 | 0 | 0 | 0 | 0 | 0 | 0 | 0 | 0 | 1 | 0 |
| hipA | 0 | 0 | 1 | 0 | 0 | 0 | 0 | 0 | 0 | 0 | 0 |
| hisC2 | 0 | 0 | 0 | 0 | 0 | 0 | 0 | 0 | 0 | 0 | 0 |
| hns | 0 | 0 | 0 | 0 | 0 | 0 | 0 | 1 | 0 | 0 | 0 |
| hsdR | 0 | 0 | 0 | 0 | 0 | 0 | 2 | 0 | 0 | 0 | 0 |
| htrC | 0 | 0 | 0 | 0 | 1 | 0 | 0 | 0 | 0 | 0 | 0 |
| hup | 0 | 0 | 0 | 0 | 0 | 0 | 2 | 1 | 0 | 0 | 0 |
| hxpA | 0 | 0 | 1 | 0 | 0 | 0 | 0 | 0 | 0 | 0 | 0 |
| ilvE | 0 | 0 | 0 | 0 | 0 | 0 | 0 | 0 | 0 | 0 | 0 |
| imm | 0 | 0 | 0 | 0 | 0 | 0 | 0 | 0 | 0 | 3 | 0 |
| invA | 2 | 0 | 0 | 0 | 0 | 0 | 0 | 0 | 0 | 0 | 0 |
| IS1 family transposase ISPda1 | 0 | 0 | 0 | 0 | 0 | 0 | 1 | 0 | 0 | 0 | 0 |
| IS110 family transposase ISCps1 | 0 | 0 | 0 | 0 | 0 | 0 | 0 | 0 | 0 | 0 | 0 |
| IS110 family transposase ISSde4 | 0 | 0 | 0 | 0 | 0 | 0 | 0 | 0 | 0 | 0 | 0 |
| IS110 family transposase ISSpi6 | 0 | 0 | 0 | 2 | 0 | 0 | 1 | 0 | 0 | 0 | 0 |
| IS200/IS605 family transposase ISEc42 | 0 | 0 | 0 | 0 | 0 | 0 | 0 | 1 | 0 | 0 | 0 |
| IS200/IS605 family transposase ISEc46 | 0 | 0 | 0 | 0 | 0 | 0 | 0 | 2 | 0 | 0 | 0 |
| IS21 family transposase ISVch3 | 0 | 0 | 0 | 1 | 0 | 0 | 0 | 0 | 0 | 0 | 0 |
| IS3 family transposase ISBps2 | 0 | 0 | 0 | 1 | 0 | 0 | 0 | 0 | 0 | 0 | 0 |
| IS3 family transposase ISThsp4 | 0 | 0 | 0 | 0 | 0 | 0 | 0 | 2 | 0 | 0 | 0 |
| IS3 family transposase ISVch4 | 0 | 0 | 0 | 0 | 0 | 0 | 4 | 0 | 0 | 0 | 0 |
| IS30 family transposase ISSde3 | 0 | 0 | 0 | 1 | 0 | 0 | 0 | 0 | 0 | 0 | 0 |
| IS4 family transposase ISAzvi5 | 0 | 0 | 0 | 0 | 0 | 0 | 1 | 0 | 0 | 0 | 0 |
| IS4 family transposase ISCro6 | 0 | 0 | 0 | 0 | 0 | 0 | 3 | 0 | 0 | 0 | 0 |
| IS4 family transposase ISPcc6 | 0 | 0 | 0 | 0 | 0 | 0 | 2 | 1 | 0 | 0 | 0 |
| IS4 family transposase ISVa18 | 0 | 0 | 0 | 0 | 0 | 0 | 3 | 0 | 0 | 0 | 0 |
| IS4 family transposase ISVsa5 | 0 | 0 | 0 | 0 | 0 | 0 | 1 | 0 | 0 | 0 | 0 |
| IS5 family transposase ISVha3 | 0 | 0 | 1 | 0 | 0 | 0 | 1 | 0 | 0 | 0 | 0 |
| IS6 family transposase ISPpr9 | 1 | 2 | 4 | 2 | 1 | 1 | 4 | 4 | 0 | 0 | 0 |
| IS630 family transposase ISVa15 | 0 | 2 | 6 | 0 | 0 | 0 | 1 | 0 | 0 | 0 | 0 |
| IS66 family transposase ISVa11 | 0 | 0 | 0 | 0 | 0 | 0 | 4 | 0 | 0 | 0 | 0 |
| IS66 family transposase ISVa5 | 0 | 0 | 0 | 0 | 0 | 0 | 0 | 1 | 0 | 0 | 0 |
| IS66 family transposase ISVa9 | 0 | 0 | 0 | 0 | 0 | 0 | 1 | 0 | 0 | 0 | 0 |
| IS91 family transposase ISShvi3 | 0 | 0 | 0 | 0 | 0 | 0 | 0 | 0 | 0 | 0 | 0 |
| IS91 family transposase ISVsa9 | 1 | 0 | 0 | 0 | 0 | 0 | 1 | 1 | 0 | 0 | 0 |
| ISAs1 family transposase ISVha1 | 0 | 0 | 0 | 0 | 0 | 0 | 1 | 0 | 0 | 0 | 0 |
| Isatin hydrolase | 0 | 0 | 0 | 0 | 0 | 0 | 0 | 0 | 0 | 0 | 0 |
| ISNCY family transposase ISLad2 | 0 | 0 | 0 | 0 | 0 | 0 | 1 | 0 | 0 | 0 | 0 |
| ISNCY family transposase ISLad2 | 0 | 0 | 0 | 0 | 0 | 0 | 0 | 0 | 0 | 0 | 1 |
| ISNCY family transposase ISVisp7 | 0 | 0 | 0 | 0 | 0 | 0 | 0 | 0 | 0 | 0 | 2 |
| kynA | 0 | 0 | 0 | 0 | 0 | 0 | 0 | 0 | 0 | 0 | 0 |
| kynu | 0 | 0 | 0 | 0 | 0 | 0 | 0 | 0 | 0 | 0 | 0 |
| lagD | 0 | 0 | 0 | 0 | 0 | 0 | 0 | 0 | 0 | 0 | 0 |
| lcdH | 0 | 0 | 0 | 0 | 0 | 0 | 0 | 0 | 0 | 0 | 0 |
| lgrD | 0 | 0 | 0 | 0 | 0 | 0 | 1 | 0 | 0 | 0 | 0 |
| lysO | 0 | 0 | 0 | 0 | 0 | 1 | 0 | 0 | 0 | 0 | 0 |
| mhpA | 0 | 0 | 0 | 0 | 0 | 0 | 0 | 0 | 0 | 0 | 0 |
| moaA | 0 | 0 | 0 | 0 | 0 | 0 | 0 | 0 | 0 | 0 | 1 |
| mtcA2 | 0 | 0 | 0 | 0 | 0 | 0 | 0 | 0 | 0 | 0 | 1 |
| N-carbamoyl-D-amino acid hydrolase | 0 | 0 | 0 | 0 | 0 | 0 | 0 | 0 | 0 | 3 | 0 |
| nimR | 0 | 0 | 0 | 0 | 0 | 0 | 0 | 0 | 0 | 0 | 0 |
| nrgA | 0 | 0 | 0 | 0 | 0 | 0 | 0 | 0 | 0 | 1 | 0 |
| ntdA | 0 | 0 | 0 | 0 | 0 | 0 | 1 | 0 | 0 | 0 | 0 |
| ntdB | 0 | 0 | 0 | 0 | 0 | 0 | 1 | 0 | 0 | 0 | 0 |
| ntdC | 0 | 0 | 0 | 0 | 0 | 0 | 1 | 0 | 0 | 0 | 0 |
| nucM | 0 | 0 | 0 | 0 | 0 | 0 | 1 | 0 | 0 | 0 | 0 |
| ompA | 0 | 0 | 0 | 0 | 0 | 0 | 0 | 0 | 0 | 0 | 0 |
| parA | 0 | 0 | 0 | 0 | 0 | 0 | 0 | 0 | 0 | 1 | 0 |
| pdeG | 0 | 0 | 1 | 0 | 0 | 0 | 0 | 0 | 0 | 0 | 0 |
| pgrR | 0 | 0 | 1 | 0 | 0 | 0 | 0 | 0 | 0 | 0 | 0 |
| pinR | 0 | 0 | 0 | 0 | 0 | 0 | 0 | 0 | 0 | 0 | 0 |
| pndA | 0 | 0 | 0 | 0 | 0 | 0 | 0 | 0 | 0 | 0 | 0 |
| pndA | 1 | 0 | 0 | 0 | 0 | 0 | 0 | 0 | 0 | 0 | 0 |
| proQ | 0 | 0 | 4 | 0 | 0 | 0 | 0 | 0 | 0 | 0 | 0 |
| prtR | 0 | 0 | 0 | 0 | 0 | 0 | 0 | 0 | 0 | 0 | 0 |
| pscF | 2 | 0 | 0 | 0 | 0 | 0 | 0 | 0 | 0 | 0 | 0 |
| ptlH | 0 | 0 | 0 | 0 | 0 | 0 | 1 | 1 | 0 | 0 | 0 |
| Putative nickel-responsive regulator | 0 | 0 | 0 | 0 | 0 | 0 | 0 | 0 | 0 | 1 | 0 |
| putative oxidoreductase | 0 | 0 | 0 | 0 | 1 | 1 | 0 | 0 | 0 | 0 | 0 |
| queuosine precursor transporter | 0 | 0 | 0 | 0 | 0 | 0 | 0 | 0 | 0 | 0 | 0 |
| ramA | 0 | 0 | 0 | 0 | 0 | 0 | 0 | 0 | 0 | 0 | 0 |
| ramC | 0 | 0 | 0 | 0 | 0 | 0 | 0 | 0 | 0 | 0 | 0 |
| rclC | 0 | 0 | 0 | 0 | 0 | 0 | 1 | 0 | 0 | 0 | 0 |
| recX | 0 | 0 | 1 | 0 | 0 | 0 | 2 | 0 | 0 | 0 | 0 |
| rep | 0 | 0 | 1 | 0 | 0 | 0 | 0 | 0 | 0 | 0 | 0 |
| repA | 0 | 2 | 0 | 0 | 0 | 0 | 1 | 0 | 1 | 1 | 0 |
| rhaR | 0 | 0 | 1 | 0 | 0 | 0 | 0 | 0 | 0 | 0 | 0 |
| rhaS | 0 | 0 | 0 | 1 | 0 | 0 | 0 | 0 | 0 | 0 | 0 |
| rhsC | 0 | 0 | 0 | 0 | 0 | 0 | 0 | 0 | 0 | 0 | 0 |
| ribD | 0 | 0 | 0 | 1 | 0 | 0 | 0 | 0 | 0 | 0 | 0 |
| ribF | 0 | 0 | 1 | 1 | 0 | 0 | 0 | 0 | 0 | 0 | 0 |
| rizA | 0 | 0 | 0 | 0 | 0 | 0 | 0 | 0 | 0 | 0 | 0 |
| rutB | 0 | 0 | 0 | 0 | 0 | 0 | 0 | 0 | 0 | 0 | 0 |
| sasA | 0 | 0 | 0 | 0 | 0 | 0 | 1 | 0 | 0 | 0 | 0 |
| sctC | 6 | 0 | 0 | 0 | 0 | 0 | 0 | 0 | 0 | 0 | 0 |
| slyA | 0 | 0 | 0 | 0 | 0 | 0 | 0 | 0 | 1 | 0 | 0 |
| smc | 0 | 0 | 0 | 0 | 0 | 0 | 0 | 0 | 1 | 0 | 0 |
| sodC | 0 | 0 | 0 | 0 | 0 | 0 | 1 | 0 | 0 | 0 | 0 |
| soj | 1 | 0 | 1 | 0 | 0 | 0 | 1 | 0 | 0 | 0 | 0 |
| soxR | 0 | 0 | 0 | 0 | 0 | 0 | 0 | 1 | 0 | 0 | 0 |
| spaR | 1 | 0 | 0 | 0 | 0 | 0 | 0 | 0 | 0 | 0 | 0 |
| tcpE | 0 | 0 | 0 | 0 | 0 | 0 | 0 | 1 | 0 | 0 | 0 |
| TDO2 | 0 | 0 | 0 | 0 | 0 | 0 | 0 | 0 | 0 | 0 | 0 |
| tetA | 0 | 0 | 0 | 0 | 0 | 0 | 0 | 0 | 0 | 0 | 0 |
| tfdR | 0 | 0 | 0 | 0 | 0 | 0 | 0 | 0 | 0 | 0 | 0 |
| tmcA | 0 | 0 | 0 | 0 | 0 | 0 | 0 | 0 | 0 | 0 | 0 |
| Tn3 family transposase ISSod9 | 0 | 0 | 0 | 0 | 0 | 0 | 2 | 0 | 0 | 0 | 0 |
| Tn3 family transposase ISXc4 | 0 | 0 | 0 | 0 | 0 | 0 | 1 | 0 | 0 | 0 | 0 |
| tnpR | 0 | 2 | 0 | 0 | 1 | 1 | 1 | 1 | 1 | 3 | 0 |
| topA | 0 | 0 | 0 | 0 | 0 | 0 | 1 | 0 | 0 | 0 | 0 |
| topB | 0 | 0 | 0 | 0 | 0 | 0 | 5 | 2 | 0 | 0 | 0 |
| traC | 1 | 0 | 0 | 0 | 0 | 0 | 3 | 3 | 0 | 0 | 0 |
| traD | 0 | 0 | 0 | 0 | 0 | 0 | 0 | 0 | 0 | 0 | 0 |
| traN | 0 | 0 | 0 | 0 | 0 | 0 | 0 | 1 | 0 | 0 | 0 |
| tyeA | 1 | 0 | 0 | 0 | 0 | 0 | 0 | 0 | 0 | 0 | 0 |
| virB | 0 | 0 | 0 | 0 | 0 | 0 | 0 | 1 | 0 | 1 | 0 |
| virB1 | 0 | 0 | 0 | 0 | 0 | 0 | 2 | 1 | 0 | 0 | 0 |
| virB4 | 0 | 0 | 0 | 0 | 0 | 0 | 1 | 0 | 0 | 0 | 0 |
| xerC | 0 | 0 | 0 | 0 | 0 | 2 | 1 | 1 | 0 | 0 | 1 |
| xerD | 0 | 0 | 0 | 0 | 0 | 0 | 0 | 0 | 0 | 0 | 0 |
| yedJ | 0 | 0 | 0 | 0 | 0 | 0 | 0 | 0 | 0 | 0 | 2 |
| yegI | 0 | 0 | 0 | 0 | 0 | 0 | 0 | 0 | 0 | 0 | 1 |
| ygiW | 0 | 0 | 0 | 0 | 0 | 0 | 1 | 0 | 0 | 0 | 0 |
| yhcG | 0 | 0 | 0 | 0 | 0 | 0 | 0 | 1 | 0 | 0 | 0 |
| yiaC | 0 | 0 | 0 | 0 | 0 | 0 | 0 | 0 | 0 | 0 | 2 |
| yigB | 0 | 0 | 0 | 1 | 0 | 0 | 0 | 0 | 0 | 0 | 0 |
| yopB | 1 | 0 | 0 | 0 | 0 | 0 | 0 | 0 | 0 | 0 | 0 |
| yopN | 1 | 0 | 0 | 0 | 0 | 0 | 0 | 0 | 0 | 0 | 0 |
| yraJ | 0 | 0 | 0 | 0 | 0 | 0 | 0 | 0 | 0 | 1 | 0 |
| yscD | 2 | 0 | 0 | 0 | 0 | 0 | 0 | 0 | 0 | 0 | 0 |
| yscG | 2 | 0 | 0 | 0 | 0 | 0 | 0 | 0 | 0 | 0 | 0 |
| yscJ | 2 | 0 | 0 | 0 | 0 | 0 | 0 | 0 | 0 | 0 | 0 |
| yscN | 1 | 0 | 0 | 0 | 0 | 0 | 0 | 0 | 0 | 0 | 0 |
| yscU | 1 | 0 | 0 | 0 | 0 | 0 | 0 | 0 | 0 | 0 | 0 |
